# Mineral phase analysis of various marine-species shells and skeletons collected in Japan: Implications for marine biominerals

**DOI:** 10.1101/2022.10.30.514443

**Authors:** Takehiro Mitsuguchi, Keiji Minakata, Kaoru Sugihara, Masanori Hiraoka, Masa-aki Yoshida, Yoko Saito-Kokubu

**Affiliations:** Tono Geoscience Center, Japan Atomic Energy Agency, Izumi-cho, Toki, Gifu 509-5102, Japan; Yabitsu Shell Museum, Arida, Wakayama 649-0316, Japan; Graduate School of Life and Environmental Sciences, University of Tsukuba, Tsukuba, Ibaraki 305-8572, Japan; Usa Marine Biological Institute, Kochi University, Inoshiri, Usa, Tosa, Kochi 781-1164, Japan; Marine Biological Science Section, Education and Research Center for Biological Resources, Faculty of Life and Environmental Science, Shimane University, Okinoshima-cho, Oki, Shimane 685-0024, Japan

**Author notes:** **Corresponding author** ^1^ T. Mitsuguchi.

## Abstract

Mineral phase analysis was performed, using X-ray diffractometry (XRD), for marine-organism shell/skeleton samples of 146–148 extant species of the following 10 phyla (18 classes) collected in Japan: Rhodophyta (Florideophyceae), Foraminifera (Globothalamea and Tubothalamea), Porifera (Hexactinellida), Cnidaria (Anthozoa and Hydrozoa), Bryozoa (Gymnolaemata), Brachiopoda (Lingulata and Rhynchonellata), Mollusca (Bivalvia, Cephalopoda, Gastropoda and Polyplacophora), Annelida (Polychaeta), Arthropoda (Cirripedia), and Echinodermata (Asteroidea, Crinoidea and Echinoidea). Some of the species were analyzed for each specific part of their shells/skeletons. Almost all the samples exhibited any of calcite, aragonite or their mixed phase, predominantly depending on their taxonomy and shell/skeletal structures. For samples containing significant amounts of calcite, the MgCO_3_ wt % of calcite has been determined from their XRD data, which ranges from ∼0 to ∼15 wt % and indicates clear inter-taxonomic differences. Low MgCO_3_ values (∼0–4 wt %) are observed for Rhynchonellata, Bivalvia, Gastropoda and Cirripedia; intermediate values (∼4–8 wt %) for Cephalopoda; high values (∼8–15 wt %) for Florideophyceae, Globothalamea, Tubothalamea, Polychaeta, Asteroidea and Crinoidea; low-to-high values for Gymnolaemata; intermediate-to-high values for Anthozoa and Echinoidea. These MgCO_3_ data show broad trends mostly consistent with general phylogenetic evolution (i.e. very similar patterns for each phylogenetic group). Distinct within-individual variability of the MgCO_3_ content is found for regular Echinoidea species (i.e. their teeth and spines have lower MgCO_3_ values than the other skeletal parts). Correlation of the MgCO_3_ content with seawater temperature is also examined/discussed for most of the above calcite-containing classes. In order to interpret our XRD-based observations of various marine-species shells/skeletons, detailed discussions are presented by comparing with previous studies and also by using knowledge of taxonomy, shell/skeletal structures, habitats, living modes and so on. The comprehensive dataset and discussions will provide useful implications for biomineralization studies.

## INTRODUCTION

An enormous number of marine species secrete shells/skeletons inside or outside of their bodies. The shell/skeleton production especially by numerous marine invertebrates plays important roles in the biogeochemical cycle and has a significant influence on the long-term global climate change. This is due to the fact that most of the invertebrate shells/skeletons are composed of caclium carbonate (CaCO_3_), which is produced by the following simplified calcification reaction: Ca^2+^ + CO_32–_ → CaCO_3_, where (i) Ca^2+^ is derived from seawater and (ii) CO_32–_ is connected with atmospheric CO_2_ gas via the reaction chain as follows: CO_2_ (g) + H_2_O (seawater) ↔ CO_2_ (aq) + H_2_O ↔ HCO_3–_ + H^+^ ↔ CO_32–_ + 2H^+^. In addition to the calcifying species, some species secrete SiO_2_·*n*H_2_O (e.g. polycystines, hexactinellids: Ehrlich, 2011; Marron et al., 2016), SrSO_4_ (acantharians: Bütschli, 1906; Odum, 1951; Brass, 1980), organically-associated Ca_5_(PO_4_)_3_F–Ca_5_(PO_4_)_3_OH solid solution (the brachiopod genus *Lingula*: Kelly et al., 1965; Iijima and Moriwaki, 1990; Williams et al., 1994) and so on.

There exist in nature three polymorphs of CaCO_3_: calcite, aragonite and vaterite in order of decreasing stability; vaterite is very unstable and rarely found in nature. Calcite and aragonite do not considerably differ in their physical/chemical properties; for example, their specific gravities are 2.71 and 2.95, respectively; their hardness in the Mohs scale are 3 and 3.5−4, respectively; their solubility products (*K*_*sp*_) in seawater at 25°C and 35‰ salinity are ∼4.4 × 10^−7^ mol^2^ kg^−2^ and ∼6.6 × 10^−7^ mol^2^ kg^−2^, respectively (Morse et al., 1980). These values are based on inorganic mineralogy/chemistry and may be slightly different from those of biogenic calcite and aragonite containing some impurities.

Inorganically formed aragonites, when heated, are transformed to calcite from ∼400°C to ∼500°C; whereas, the transformation of marine biogenic aragonites (coral skeletons and bivalve shells) to calcite occurs from ∼280°C to ∼380°C (Yoshioka and Kitano, 1985; Antao and Hassan, 2010). From the viewpoint of the pressure-temperature phase diagram of CaCO_3_, calcite is clearly more stable than aragonite in the present condition of the Earth’s surface. On the other hand, many marine species secrete aragonite (e.g. scleractinian corals, chitons, most bivalves and gastropods), some secrete calcite (e.g. sea urchins, starfish and barnacles), and others secrete both aragonite and calcite (e.g. some bivalves, gastropods and calcareous tubeworms). It has been demonstrated, in natural observations and laboratory experiments, that aragonite is preferentially formed from the present seawater with the Mg/Ca mole ratio of ∼5.2, and that the aqueous Mg/Ca ratio is the dominant inorganic factor controlling the selective formation of aragonite and/or calcite. The present-seawater Mg/Ca mole ratio has a very small or negligible spatial (inter-regional) variability (presumably ≤ 1 % in most open oceans: e.g. Tsunogai et al., 1968, 1973), because both Mg and Ca are major dissolved elements (i.e. conservative elements) in the present ocean. An inorganic precipitation experiment of Balthasar and Cusack (2015) provides the following observations: (i) aragonite (> 98 %) is formed in a 25°C CaCO_3_-supersaturated seawater-like solution with the Mg/Ca mole ratio of > 2; (ii) both aragonite and calcite are evidently formed in the solution with the Mg/Ca ratio of 0.5–2; (iii) calcite (> 98 %) is formed in the solution with the Mg/Ca ratio of < 0.5; (iv) the solution Mg/Ca ratios under which aragonite is exclusively or significantly formed decrease with increasing the solution temperature. These results have been demonstrated *in vivo* by Higuchi et al. (2017); they observed that aragonite-secreting scleractinian corals produced significant amounts of calcite when cultivated in artificial seawaters with Mg/Ca mole ratios of 0.5 and 1.0. On the other hand, the above-mentioned fact that some marine species secrete shells/skeletons of calcite only or both calcite and aragonite in the present seawater suggests significant biologial effects on the inorganic CaCO_3_ preicpitation. It has been shown or suggested for various CaCO_3_-secreting marine species (e.g. pearl oysters: *Pinctada*, mussels: *Mytilus*, abalones: *Haliotis*, coccolithophores: *Coccolithus*, and foraminifera: *Amphistegina*) that *in vivo*-synthesized specific organic compounds (especially proteins) play dominant roles in the selective formation of aragonite and/or calcite (e.g. Watabe and Wilbur, 1960; Belcher et al., 1996; Falini et al., 1996; Tsukamoto et al., 2004; Takeuchi et al., 2008; Suzuki et al., 2009; Marie et al., 2012; Durak et al., 2017; Tyszka et al., 2019). Checa et al. (2007) observed that some bivalve species with predominantly calcite shells secreted a considerable amount of aragonite when cultivated in artificial seawater with abnormally high Mg/Ca mole ratios of 8.3–9.2, which may indicate that, in the normal seawater condition, their biological effects exceed or inhibit the effect of seawater Mg/Ca ratio.

It has been estimated that, through the Phanerozoic eon (541 million years ago to present), the seawater Mg/Ca mole ratio varied drastically between ∼1 and ∼5.2; the estimation is not only derived from a geochemical mixing model concerning the mid-ocean ridge hydrothermal flux and river water input (Hardie, 1996; Demicco et al. 2005) but also generally consistent with paleoseawater Mg/Ca variations reconstructed from (i) chemical composition of fluid inclusions in marine halite (e.g. Lowenstein et al., 2001; Horita et al., 2002; Brennan and Lowenstein, 2002; Siemann, 2003; Brennan et al., 2004; Timofeeff et al., 2006) and (ii) Mg content (or Mg/Ca ratio) of well-preserved calcite skeletons of fossil echinoderms (echinoids and crinoids: Dickson, 2002, 2004). Combined with many inorganic precipitation experiments such as performed by Balthasar and Cusack (2015) (see above), it has been deduced that, in the Phanerozoic ocean, aragonite (calcite) tended to be preferentially formed or secreted during periods of the seawater Mg/Ca mole ratio higher (lower) than ∼2 (e.g. Hardie, 1996; Stanley and Hardie, 1998; Ries, 2010; Lowenstein et al., 2014).

It has been observed, in artificial seawater with the Mg/Ca mole ratio of ∼1.0–6.7, that echinoids secrete calcite skeletons only, and that the Mg/Ca ratio of their skeletons is positively correlated with both the ambient-seawater Mg/Ca ratio and temperature (e.g. Ries, 2004, 2010). The positive correlation of the calcite Mg/Ca ratio with temperature has been observed in various marine-species shells/skeletons (e.g. bivalves, sea urchins, foraminifera, coralline algae, ostracods, crinoids and deep-sea octocorals: Chave, 1954; Weber, 1973; Nürnberg et al., 1996; Ries, 2004, 2010; Freitas et al., 2005; Yoshimura et al., 2011) and also in inorganic precipitation experiments (e.g. Kitano and Kanamori, 1966; Katz, 1973; Oomori et al., 1987; Mucci, 1987); these previously-observed Mg/Ca–temperature relationships exhibit significant differences between taxa or experimental conditions. It has also been observed that the Mg content of marine biogenic calcites affects their physical/chemical properties; for example, (i) the hardness of marine-species calcitic shells/skeletons increases with their Mg content (e.g. bivalves: Kunitake et al., 2012; echinoids: Long et al., 2014) and (ii) marine biogenic calcites with higher Mg content (≥ ∼8 wt % MgCO_3_) are clearly more soluble in seawater than those with lower Mg content (≤ ∼4 wt % MgCO_3_) (e.g. Morse and Mackenzie, 1990; Railsback, 2006). Mg^2+^ ions, when incorporated into calcite, replace Ca^2+^ ions in its crystal lattice; therefore, both the expressions ‘Mg/Ca mole ratio’ and ‘MgCO_3_ wt %’ are equivalent to the Mg concentration in calcite.

It can be considered that the drastic variation of seawater composition over the Phanerozoic eon greatly influenced the evolution of all marine species and induced the today’s diversity of marine-species biomineralization mechanisms, including (i) the selective formation of aragonite and/or calcite and (ii) the uptake of minor elements into biominerals. Investigating crystal forms, structures and chemical compositions of marine biominerals for a wide variety of taxa would provide important information for taxonomic characteristics of biomineralization mechanisms. In the current study, we performed X-ray diffraction (XRD) analyses of shell/skeleton samples of 146–148 extant marine species of 10 phyla (18 classes) collected from various locations in Japan, in order to reveal their mineral phases and determine the MgCO_3_ percentage in calcite that was detected as a single or significant-amount mineral phase in many of the samples. Some of the species were analyzed for each specific part of their shells/skeletons, while others for the whole shells/skeletons. The mineral-phase and MgCO_3_ data were investigated concerning inter-taxonomic and/or within-individual differences that might be indicative of differences in biomineralization processes. Further, the MgCO_3_ data (i.e. Mg concentrations in calcite) of near-sea-surface species were examined for correlation with sea-surface temperatures (SSTs) of their habitat regions. This comprehensive research will provide important implications for marine biomineralization studies and other research fields such as ecology, biogeochemistry and paleontology. These kinds of marine biomineral data in Japan have been documented in scattered publications and scarcely compiled, while some compilations have been published for species collected outside Japan (e.g. Lowenstam and Weiner, 1989).

## MATERIALS AND METHODS

Shell/skeleton-secreting extant marine organisms of 146–148 species (172 individuals) of 10 phyla (including 18 classes) were collected in the period of 1993–2021 from various locations in Japan (Fig. 1), with the exception that 3 species of them were collected in the 1970s–1980s. The 10 phyla (18 classes) are as follows: Annelida (Polychaeta), Arthropoda (Cirripedia), Brachiopoda (Lingulata and Rhynchonellata), Bryozoa (Gymnolaemata), Cnidaria (Anthozoa and Hydrozoa), Echinodermata (Asteroidea, Crinoidea and Echinoidea), Foraminifera (Globothalamea and Tubothalamea), Mollusca (Bivalvia, Cephalopoda, Gastropoda and Polyplacophora), Porifera (Hexactinellida), and Rhodophyta (Florideophyceae). All of the species were alive when collected, except for 6 species being dead with intact shells/skeletons and no organic tissues. All the collected species were dried in air for a few days or more. For the live-collected species of Cirripedia, Lingulata, Rhynchonellata, Bivalvia, Cephalopoda and Gastropoda, organic tissues were removed from the dried specimens, using knives, tweezers and cleaning brushes, to obtain shell/skeleton samples. For the live-collected species of Polychaeta, Anthozoan soft coral, Asteroidea, Crinoidea, Echinoidea, Polyplacophora and Florideophyceae, organic tissues were decomposed and removed by immersing the dried specimens into a commercial liquid bleach with 5 % NaClO, 0.5 % NaOH and some surfactant for a few or several days, resulting in remaining shells/skeletons. Neither of these treatments for removing organic tissues were applied to the other live-collected species of Gymnolaemata, Anthozoan hard coral, coral-like Hydrozoa, Globothalamea, Tubothalamea and Hexactinellida, because they had very high proportions of shells/skeletons relative to organic tissues and because three of the hard coral species appeared to entirely or partly consist of hard organic skeletons.

**Fig. 1.**
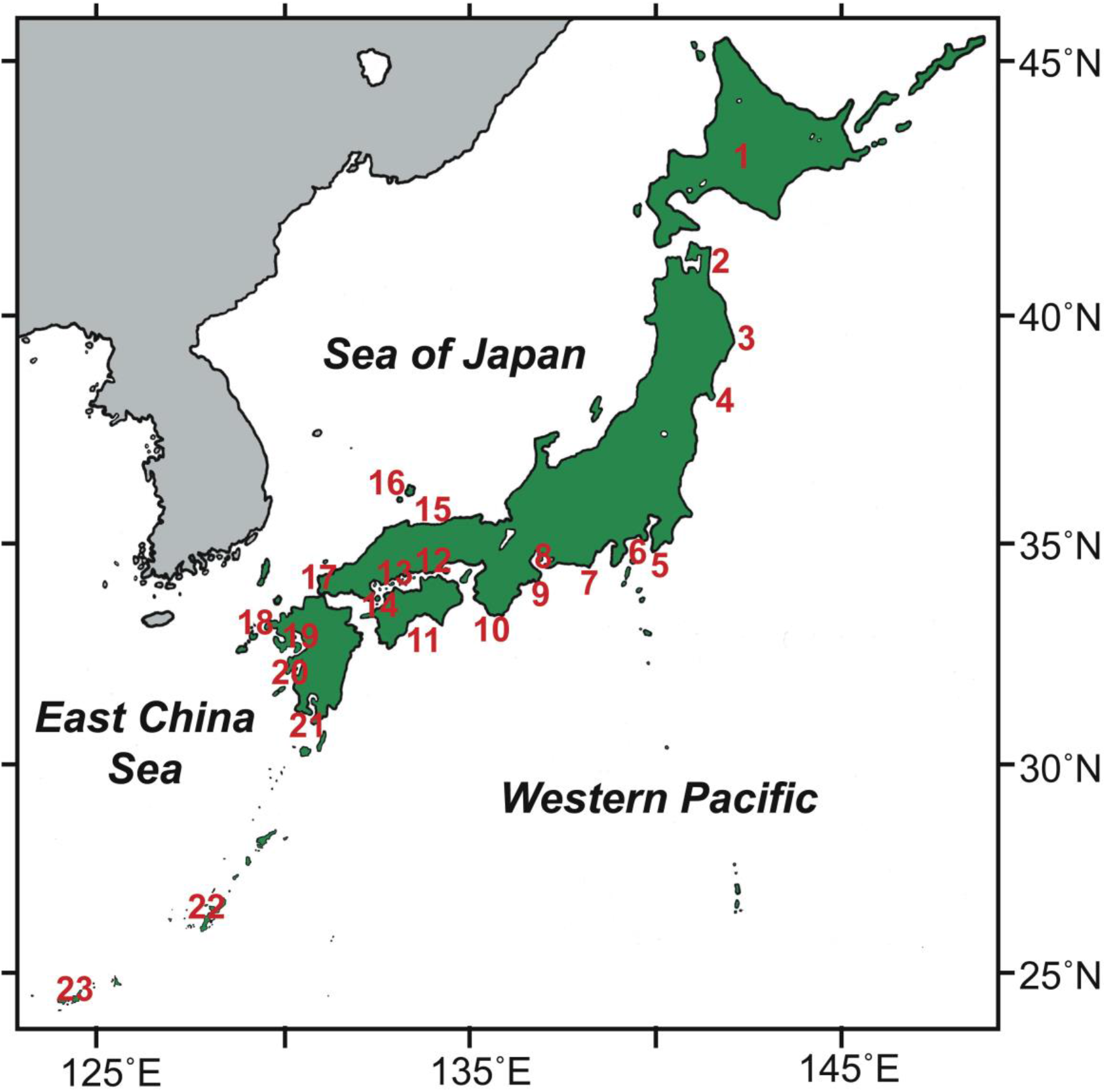
Map of Japanese islands showing locations (1–23) of sample collection for extant marine-species shells/skeletons analyzed in this study. The location numbers correspond to those shown in Tables 1 and 2.

The whole or specific part(s) of each species shell/skeleton sample were cleaned with Milli-Q water in an ultrasonic bath, dried in air, and then ground into powder for XRD analysis to determine their mineral phases; here, it should be noted that all the species used in this study were adults with well-developed shells/skeletons. Prior to the Milli-Q-water cleaning step, each sample was checked for foreign substances and biofouling (i.e. adhesion and contamination of other species’ shells/skeletons), which were removed if any by using tweezers and/or a handpiece grinder. For the species of Polychaeta, Rhynchonellata, Gymnolaemata, Anthozoa, Hydrozoa, Asteroidea, Globothalamea, Tubothalamea, Cephalopoda, Hexactinellida, and Florideophyceae, the whole of each species shell/skeleton sample was ground into powder by using an agate mortar and pestle (AMP) or exceptionally by using another method for three samples (see below). In the case of the Anthozoan hard-coral and Hydrozoa species, a skeleton sample containing all morphological structures, which was obtained from each specimen using a handpiece and diamond cutting discs, was subjected to the above-mentioned process of Milli-Q-water cleaning to AMP powderization, except that three Anthozoan skeleton samples apparently consisting of hard organic material were ground into powder by using a diamond file. For the species of Cirripedia, Crinoidea, Echinoidea, Bivalvia, Gastropoda, and Polyplacophora, the whole or specific part(s) of each species shell/skeleton sample were ground into powder using AMP. In the case of the Bivalvia species, some specific parts (e.g. inner nacreous layer, outer prismatic layer, ventral part, dorsal/umbo part, hinge part, adductor muscle scar) or a dorsal/umbo-to-ventral part including the adductor muscle scar (assumed to be equivalent to the whole shell) were cut from each species sample (by using a handpiece and diamond cutting/grinding tools) and then subjected to the above-mentioned process of Milli-Q-water cleaning to AMP powderization. Also in the case of the Gastropoda species, some specific parts (e.g. inner nacreous layer, outer thin layer, columella, operculum, protoconch) or a longitudinal cut through the shell (assumed to be equivalent to the whole shell) were obtained from each species sample (by using a handpiece and diamond cutting/grinding tools) and then subjected to the above-mentioned process of Milli-Q-water cleaning to AMP powderization. Some of the Bivalvia and Gastropoda adult shells were small (< ∼20 mm); for these species, the whole of each shell was used for the powderization. As for three Cirripedia, one Crinoidea, and three Echinoidea species, almost all specific skeletal parts of each were separated and individually subjected to the above-mentioned process of Milli-Q-water cleaning to AMP powderization (e.g. carina, carinal latus, latus, rostrum, scutum and tergum from the Cirripedia species; stalks, cirii, arms and pinnules from the Crinoidea species; spines, test plates, teeth, hemipyramids, epiphyses and rotulae from the Echinoidea species). As for one Lingulata species, it was impossible to grind its shell sample into powder by using AMP, because of very flexible nature of the shell. The flexibility is caused by relatively high organic content of the shell (Kelly et al., 1965; Iijima and Moriwaki, 1990; Williams et al., 1994); thus, the shell samples of this species were further treated in three ways (i) with the above-mentioned commerial liquid bleach for 2 days or (ii) with 30 % H_2_O_2_ aqueous solution for 2 days (both for decomposition of the shell’s organic matter), cleaned with Milli-Q water in an ultrasonic bath, dried in air, and then successfully ground into powder using AMP or (iii) with liquid nitrogen (–196°C) and cryogenically ground into powder using AMP.

The shell/skeleton powder samples obtained from the investigated marine species were analyzed for their mineral phases, using a high-precision X-ray diffractometer (Rigaku Ultima IV) with an output stability of 0.05 %, in Tono Geoscience Center of Japan Atomic Energy Agency (JAEA); the diffractometer was operated using Cu Kα radiation (40 kV/30 mA) and a Kβ filter and employing the following scanning conditions: a 2θ range from 3° to 70°, a scan rate of 1° (2θ) per minute, a step size of 0.01° (2θ), divergence slits of 0.5° (2θ) and 10 mm (for restriction of the longitudinal width), a scattering slit of 8 mm, and a fully opened receiving slit, where a non-reflective sample holder, an adjustable knife edge and a highly-advanced X-ray detector (D/teX Ultra) were used in order to reduce the background noise and to obtain more precise and accurate data.

As shown in the Results and Discussion section below, almost all of the analyzed shells/skeletons were composed of any of aragonite, calcite or their mixed phases. Aragonite and calcite percentages of the two-phase-mixed shells/skeletons were estimated by using XRD data of standard powders prepared by mixing pure aragonite powder and pure calcite powder in various weight ratios; in other words, the relationship between the aragonite/calcite weight ratio and the aragonite/calcite diffraction-peak height ratio of the standard powders was applied to the two-phase-mixed samples. This method has been frequently used in previous studies of calcareous marine-species shells/skeletons (e.g. Chave, 1954; Davies and Hooper, 1963; Smith et al., 1998; Gray and Smith, 2004; Smith and Clark, 2010). In the current study, the method carried an uncertainty of ± 3–5 wt % in the range of ∼10–90 wt % (∼90–10 wt %) of aragonite (calcite), and detection limits of aragonite and calcite in their mixture were estimated to be ∼2 wt % and ∼0.5 wt %, respectively. For samples with significant amounts of calcite (i.e. ∼10–100 wt %), the MgCO_3_ wt % was determined using the linear relationship between the XRD-derived *d*_104_ spacing value and MgCO_3_ content in calcite (Goldsmith et al., 1961) accurately applicable within a range of 0 to ∼25 wt % MgCO_3_ (Zhang et al., 2010). This kind of XRD-based MgCO_3_ measurements for calcite samples was first reported by Chave (1952), validated by subsequent studies (e.g. Goldsmith et al., 1961; Mackenzie et al., 1983), and applied to various marine-species shells/skeletons (e.g. Chave, 1954; Rucker and Carver, 1969; Medaković et al., 1995; Smith et al., 1998; Wejnert and Smith, 2008; Taylor et al., 2009; Smith and Clark, 2010; Loxton et al., 2014, 2018; Williamson et al., 2014; Krzeminska et al., 2016). In the current study, the reproducibility of our MgCO_3_ measurements was estimated to be within 0.5 wt % by replicate measurements of some calcitic marine-species powder samples with ∼0.5 to ∼14 wt % MgCO_3_.

## RESULTS AND DISCUSSION

The XRD results for the shell/skeleton samples of 146–148 extant marine species (172 individuals) are summarized in Table 1, where the phyla are arranged in order according to the general phylogenetic tree (e.g. Hasegawa, 2017). The species/genus names and their taxonomic classification are based on the World Register of Marine Species (WoRMS), which is updated regularly (Appeltans et al., 2011). As expected, almost all the analyzed species exhibit any of aragonite, calcite or their mixed phases; other mineral or non-mineral phases are found only in 5 species. The calcite phases detected in significant amounts are categorized, on the basis of the XRD-based MgCO_3_ measurements (see Table 1), into (i) low magnesian calcite (LMC: 0–4 wt % MgCO_3_), (ii) intermediate magnesian calcite (IMC: 4–8 wt % MgCO_3_), and (iii) high magnesian calcite (HMC: >8 wt % MgCO_3_); the LMC/IMC/HMC boundaries have been adopted in many previous studies (e.g. Rucker and Carver, 1969; Smith et al., 2006). Table 2 lists regional annual-mean SSTs corresponding to all the sample locations (see Fig. 1), which are based on instrumental observations through the period 1991–2020 by Japan Meteorological Agency or local fisheries experimental stations. The SST data are used in our discussion and assessment below concerning their relation to our MgCO_3_ data of near-sea-surface species (≤ 30 m in water depth: see Table 1).

**Table 1.**
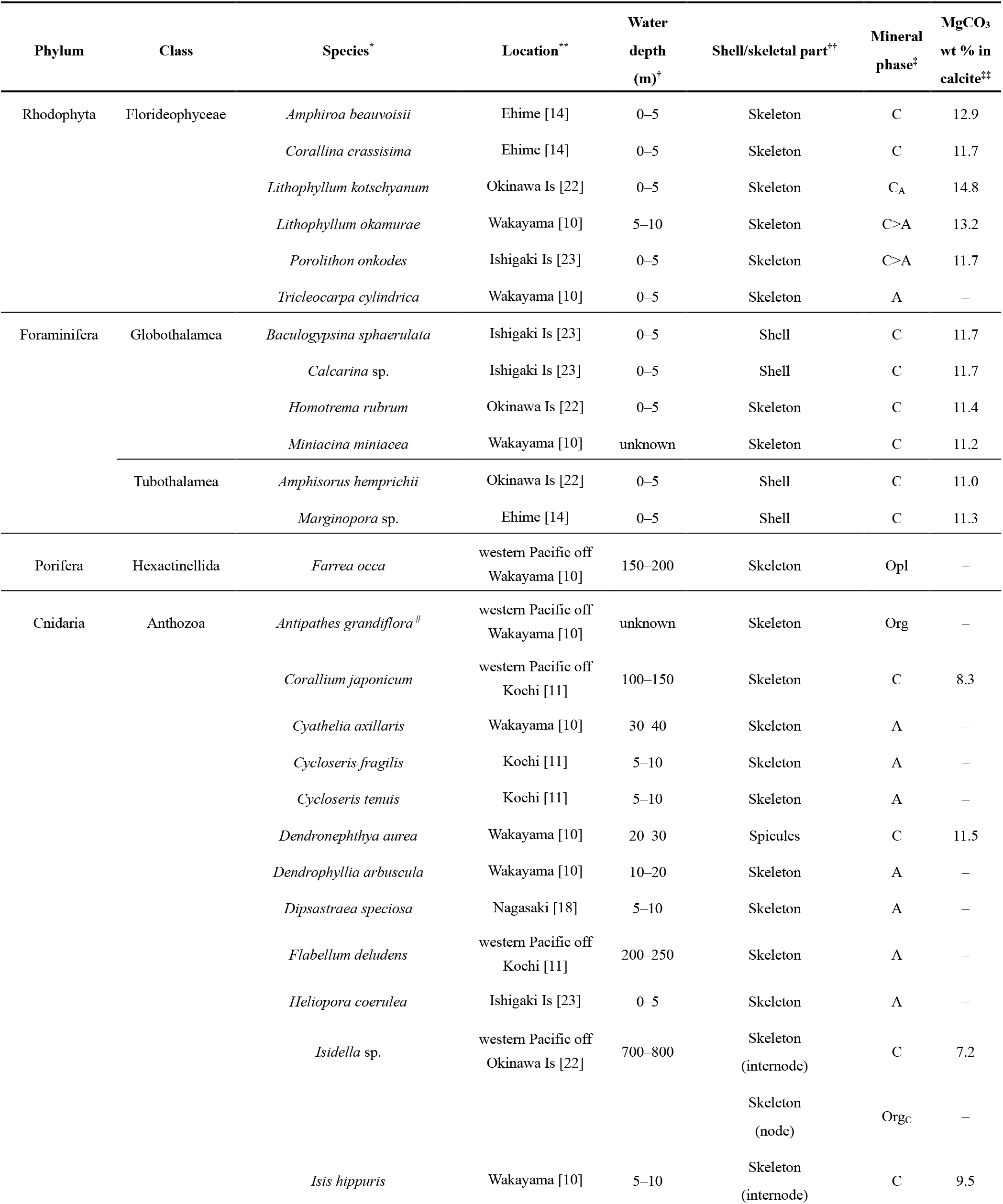
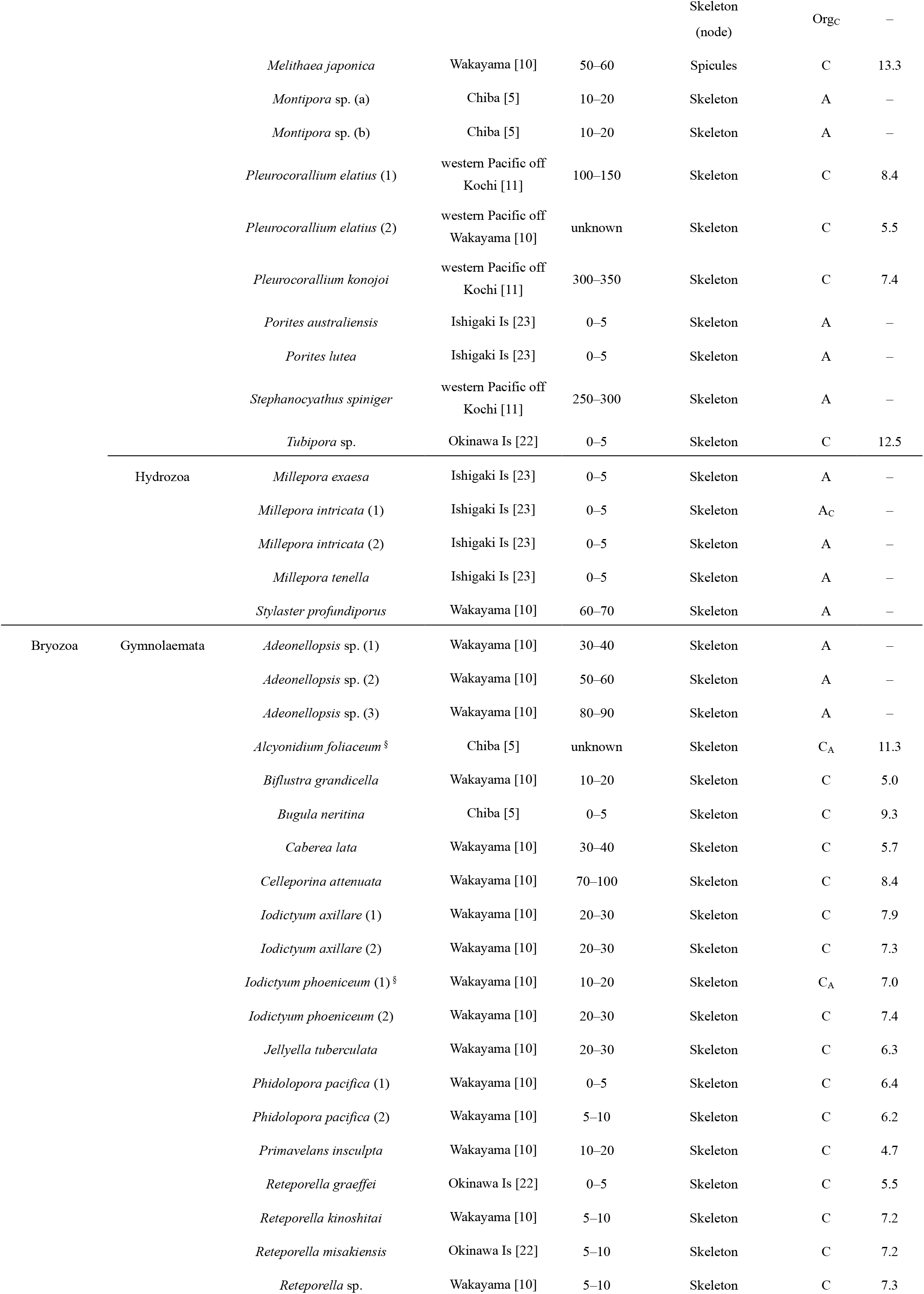
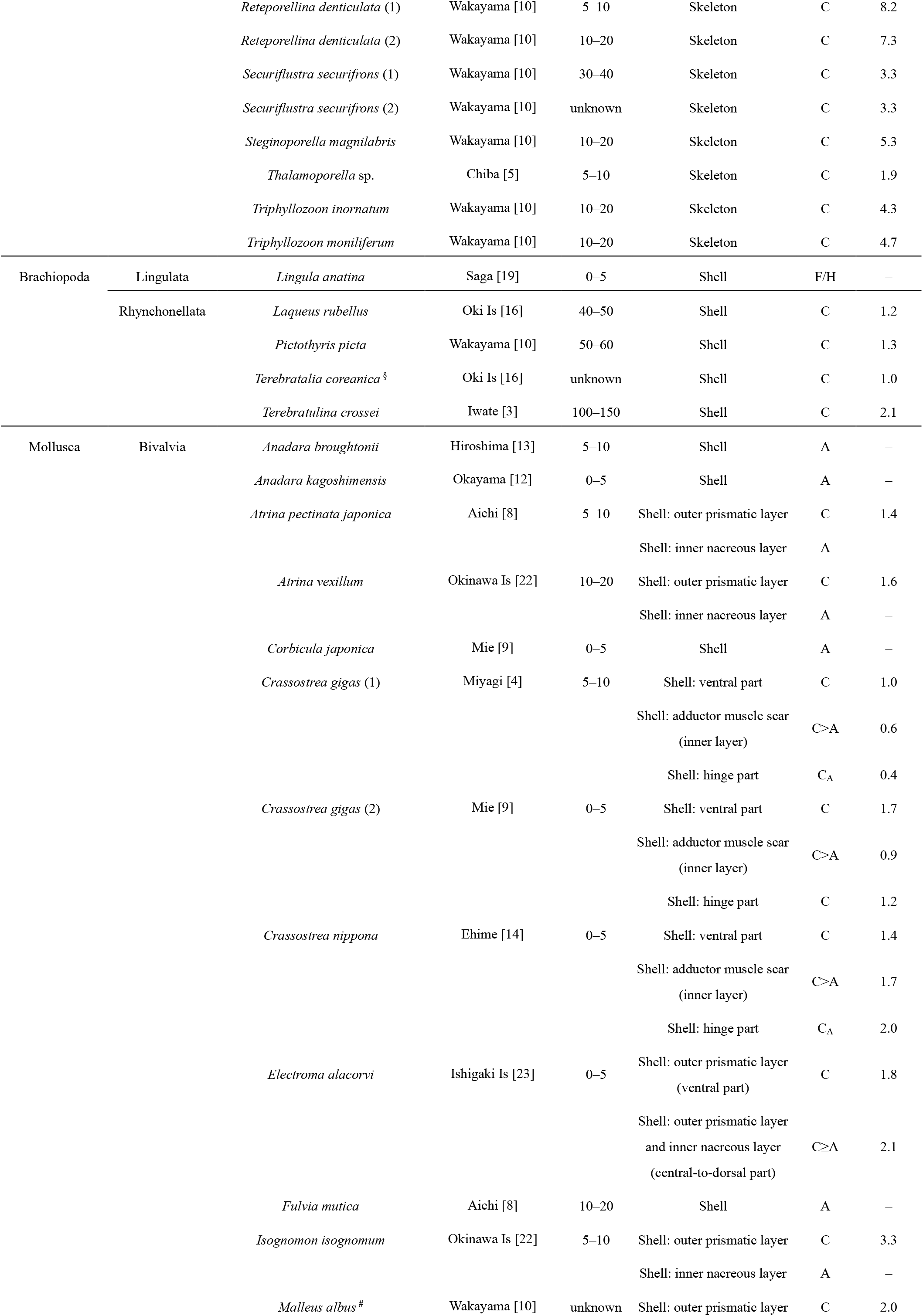
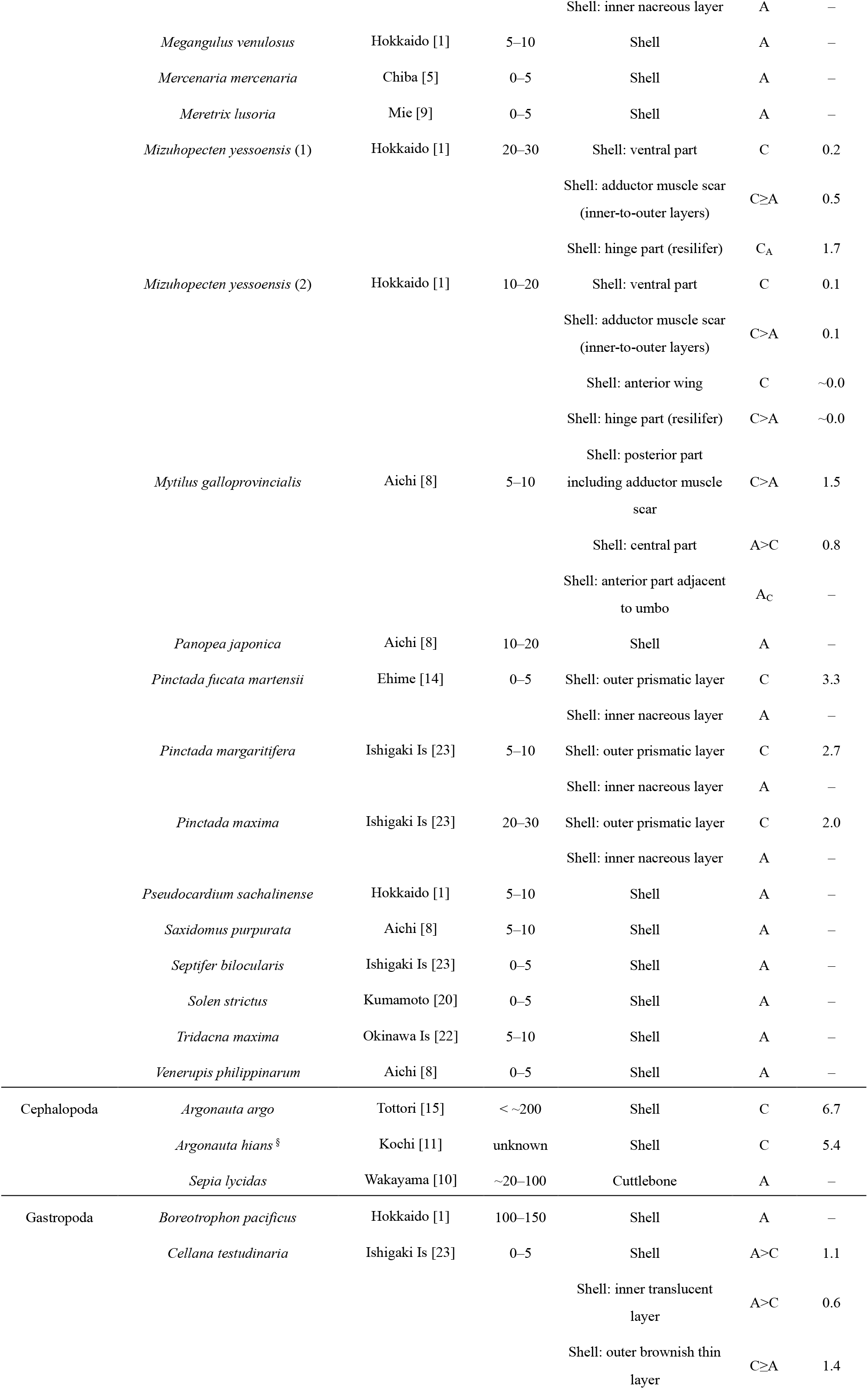
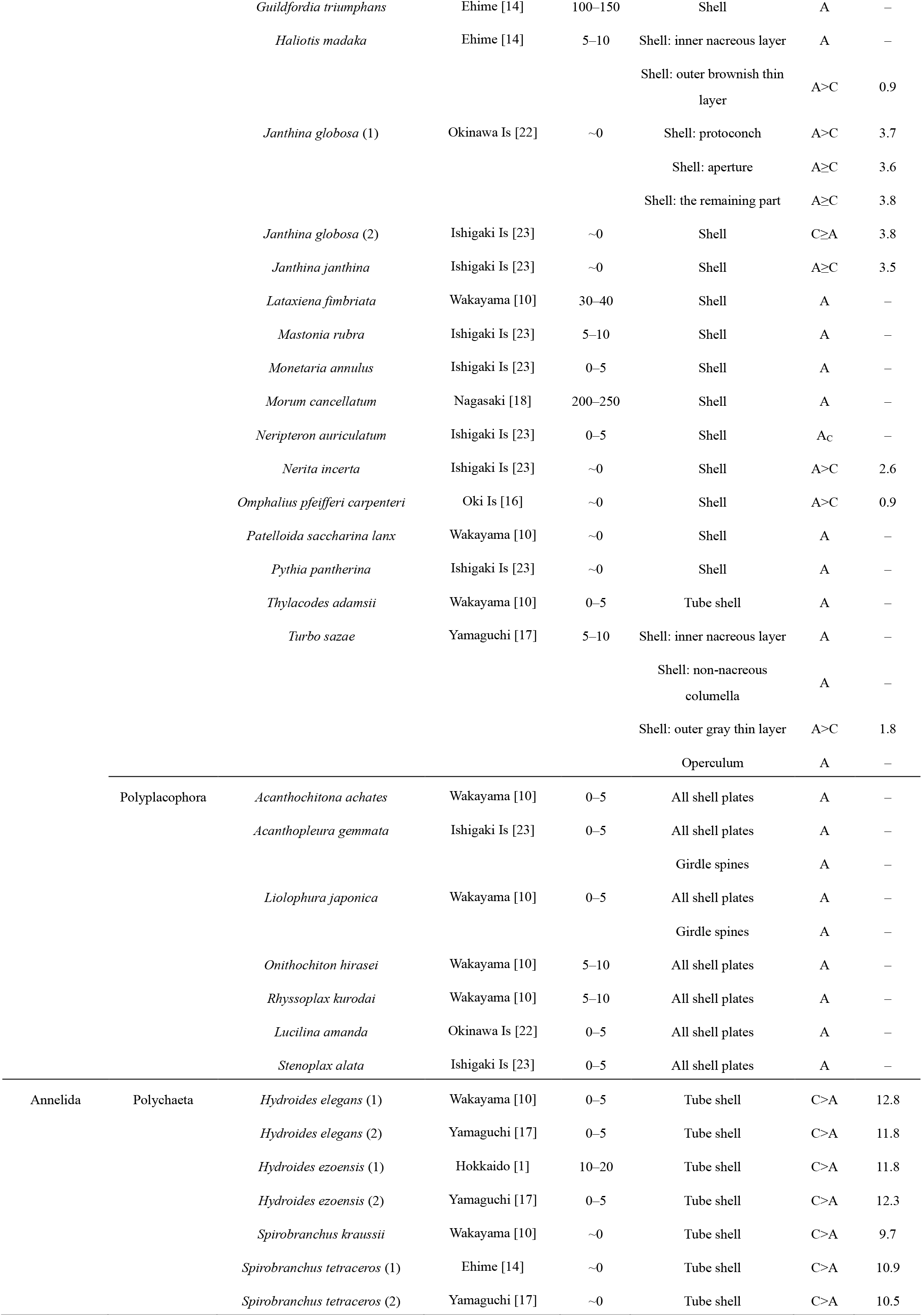
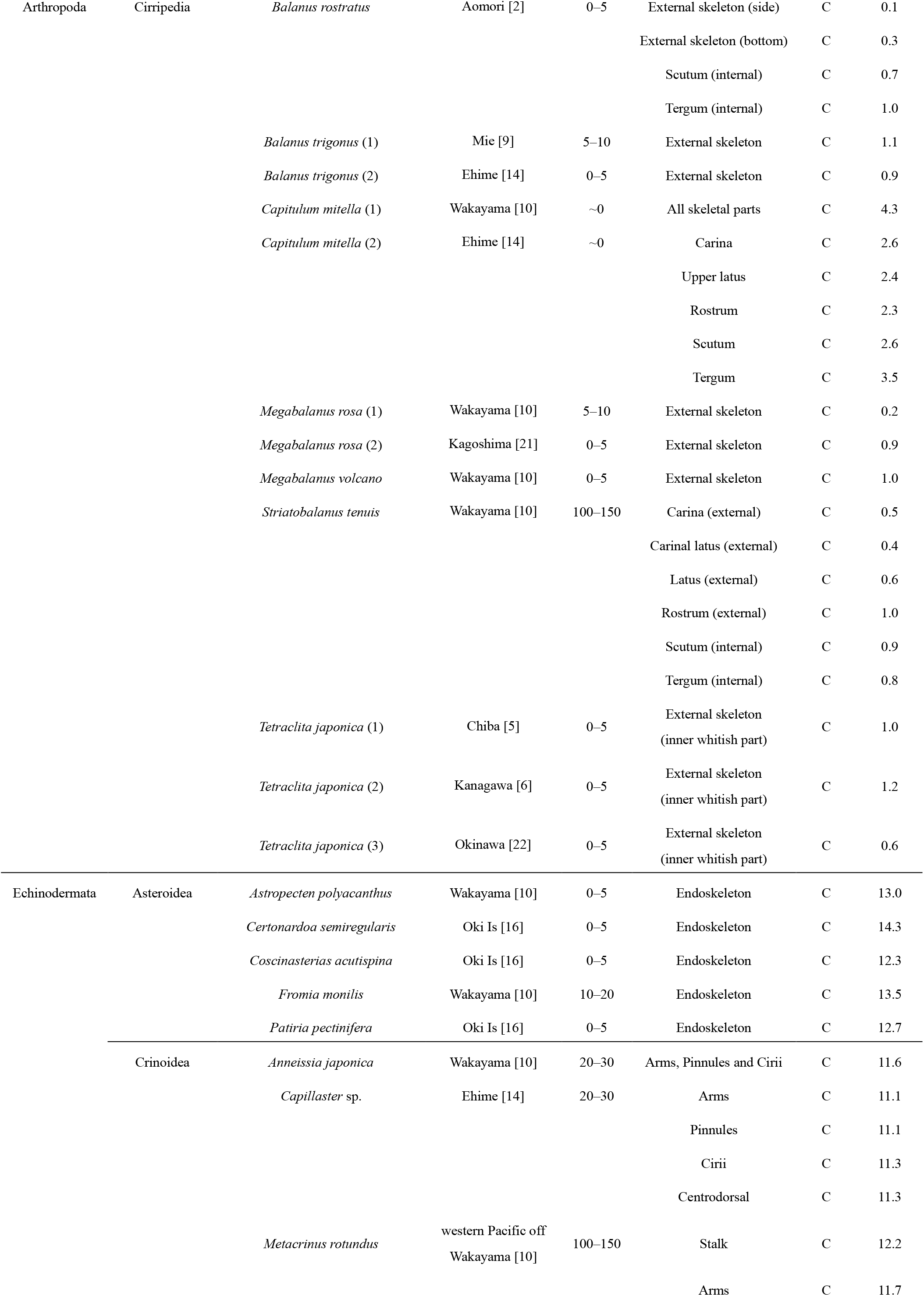
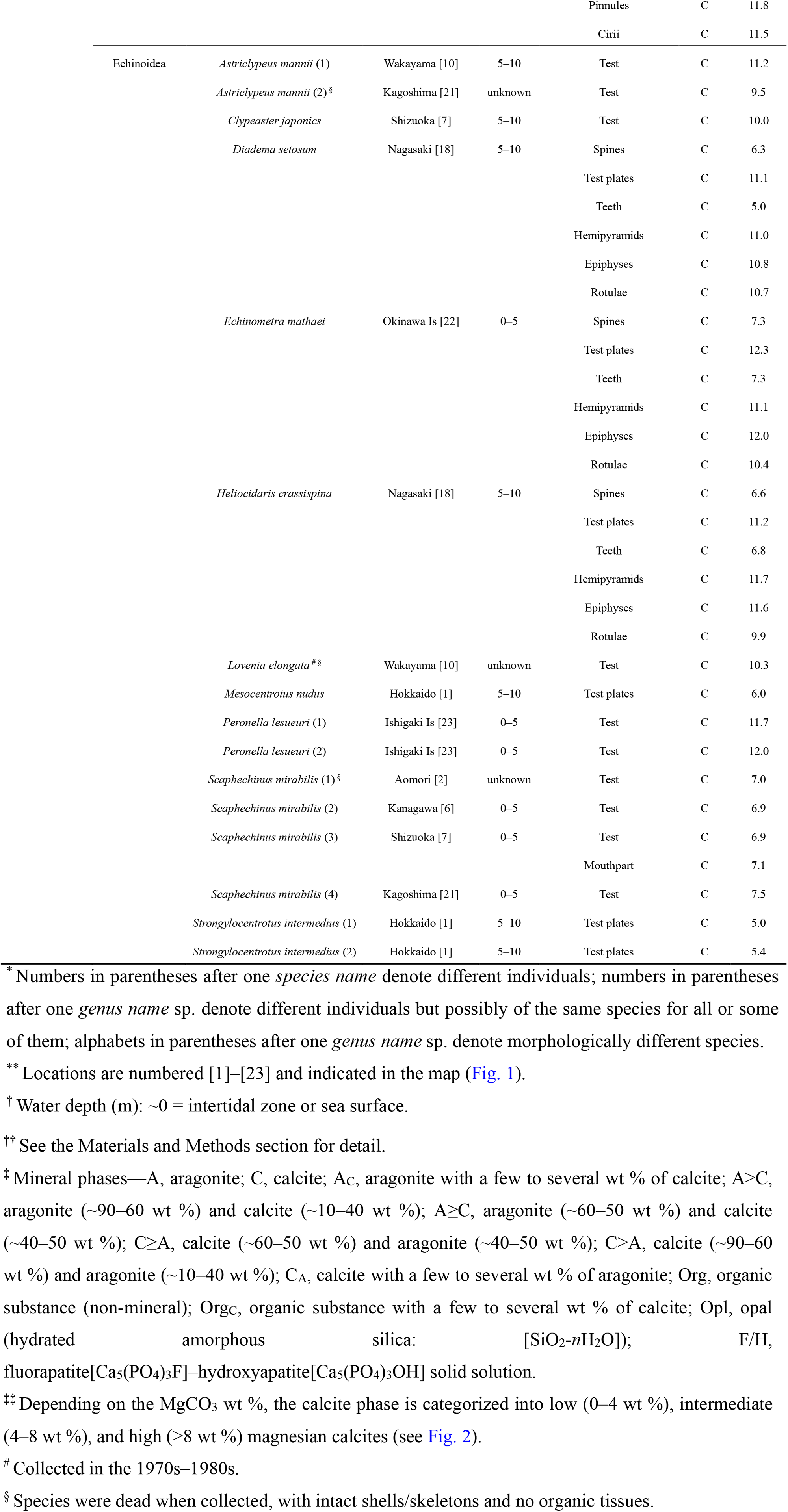
XRD-based mineral phases of marine-organism shell/skeleton samples of 146–148 extant species (172 individuals) of 10 phyla (18 classes) from various locations in Japan (Fig. 1). The phyla are arranged in order according to the general phylogenetic tree (e.g. Hasegawa, 2017). All the species were collected in 1993–2021, except for 3 species in the 1970s–1980s. For samples with significant amounts of calcite, the MgCO_3_ wt % in calcite has been determined using the XRD data.

**Table 2.** Annual-mean sea-surface temperatures (SSTs) of 23 locations in Japan where marine-organism shell/skeleton samples were collected for XRD analysis (Fig. 1). The annual-mean data are based on instrumental observations through the period 1991–2020 by Japan Meteorological Agency or local fisheries experimental stations. The annual-mean SSTs of the 13 locations [5]–[9] and [12]–[19] are not largely different, with their mean and SD of 19.6 ± 1.2°C.

| Location* | Annual mean SST (°C) |
| --- | --- |
| Hokkaido [1] | 7.2–12.7 |
| Aomori [2] | 13.7 |
| Iwate [3] | 14.3 |
| Miyagi [4] | 15.5 |
| Chiba [5] | 20.5 |
| Kanagawa [6] | 20.4 |
| Shizuoka [7] | 20.8 |
| Aichi [8] | 18.4 |
| Mie [9] | 21.3 |
| Wakayama [10] | 22.1 |
| Kochi [11] | 22.7 |
| Okayama [12] | 18.1 |
| Hiroshima [13] | 18.3 |
| Ehime [14] | 20.6 |
| Tottori [15] | 18.7 |
| Oki Is [16] | 18.6 |
| Yamaguchi [17] | 19.6 |
| Nagasaki [18] | 20.8 |
| Saga [19] | 18.5 |
| Kumamoto [20] | 21.7 |
| Kagoshima [21] | 23.6 |
| Okinawa Is [22] | 25.3 |
| Ishigaki Is [23] | 26.3 |

\* Location numbers [1]–[23] correspond to those indicated in Fig. 1 and Table 1.

### Inter-taxonomic differences in marine biomineral

Only calcite is detected in the samples of Foraminifera (Globothalamea: 4 species; Tubothalamea: 2 species), Arthropoda (Cirripedia: 7 species) and Echinodermata (Asteroidea: 5 species; Crinoidea: 3 species; Echinoidea: 10 species). The samples of Bryozoa (Gymnolaemata: 21–23 species) are constituted of calcite rarely accompanied by a trace amount (∼2–5 wt %) of aragonite, except for *Adeonellopsis* samples entirely composed of aragonite. The samples of Brachiopoda consist of calcite (Rhynchonellata: 4 species) and fluorapatite[Ca_5_(PO_4_)_3_F]–hydroxyapatite[Ca_5_(PO_4_)_3_OH] solid solution (Lingulata: 1 species). The samples of Rhodophyta (Florideophyceae: 6 species) exhibit any of calcite, aragonite or their mixture. The samples of Cnidaria (Anthozoa: 21 species; Hydrozoa: 4 species) mostly consist of either aragonite or calcite; 3 Anthozoa species have skeletons entirely or partly composed of hard organic substance. The samples of Mollusca (Bivalvia: 26 species; Cephalopoda: 3 species; Gastropoda: 17 species; Polyplacophora: 7 species) are either aragonite or aragonite-calcite mixture, except for 2 Cephalopoda species of the genus *Argonauta* entirely calcite. All the samples of Annelida (Polychaeta: 4 species) have mixture phases of calcite (∼85–65 wt %) and aragonite (∼15–35 wt %). The sample of Porifera (Hexactinellida: 1 species) is constituted of hydrated amorphous silica (i.e. opal: [SiO_2_·*n*H_2_O]).

A plot of the calcite-phase MgCO_3_ wt % versus biological classification is given in Fig. 2, where the phyla are arranged in order according to the general phylogenetic tree; this plot shows distinct inter-taxonomic differences. Calcite phases of the Rhodophyta (Florideophyceae), Foraminifera (Globothalamea and Tubothalamea), Cnidaria (Anthozoa), Annelida (Polychaeta) and Echinodermata (Asteroidea, Crinoidea and Echinoidea) species are all or mostly HMC; some of the Anthozoa and Echinoidea species have skeletons of entirely or partly IMC (e.g. teeth and spines of 3 Echinoidea species: see Table 1). On the other hand, calcite phases of the Brachiopoda (Rhynchonellata), Mollusca (Bivalvia, Cephalopoda and Gastropoda) and Arthropoda (Cirripedia) species are almost all LMC, with the exceptions of Cephalopoda *Argonauta* species being IMC. A wide range of the MgCO_3_ wt % from LMC to HMC is found for the Bryozoa (Gymnolaemata) species. As highlighted in Fig. 2, the MgCO_3_ data exhibit broad trends mostly consistent with the general phylogenetic evolution, varying from HMC/IMC of Rhodophyta, Foraminifera, Cnidaria and Bryozoa to IMC/LMC of Bryozoa, Brachiopoda, Mollusca and Arthropoda, and then back to IMC/HMC of Echinodermata; this trend does not apply to HMC of Annelida. All or some of the HMC species of Rhodophyta, Foraminifera and Cnidaria are photosynthetic or photosymbiotic (see Fig. 2); we speculate that photosynthetic activities within calcite-secreting marine species tend to significantly enhance the Mg uptake into their calcitic shells/skeletons. This speculation is in accord with a tendency of the Mg content in benthic Foraminifera species with and without symbiotic algae (see the below discussion on Foraminifera). Giving further discussion/explanation for the overall taxonomic trend in the MgCO_3_ data is, however, an issue too big/complicated for the current study, but could be done in future studies.

**Fig. 2.**
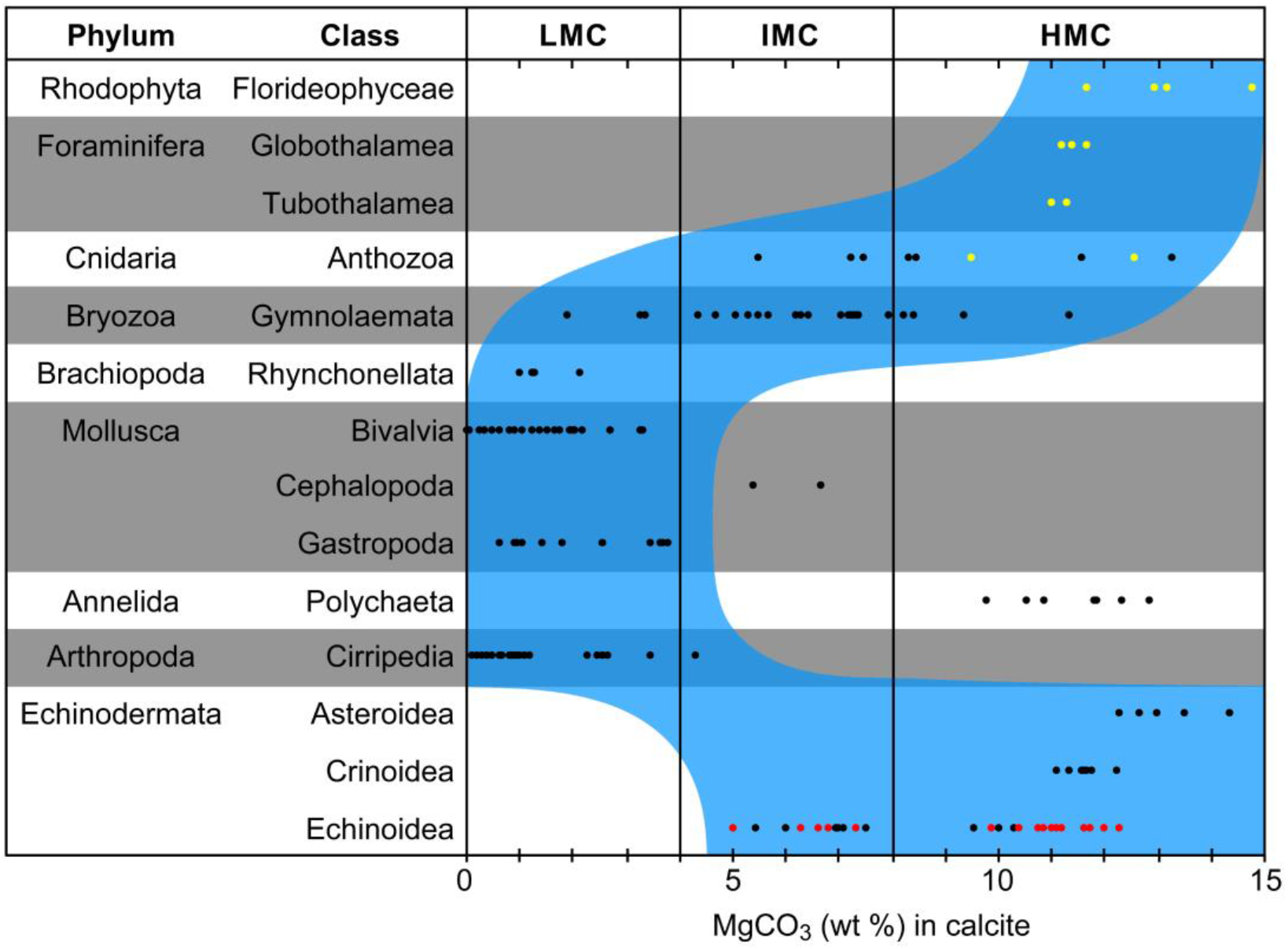
Plot of the calcite-phase MgCO_3_ wt % of extant marine-species shells/skeletons collected in Japan versus biological classifications (Table 1). The phyla are arranged in order according to the general phylogenetic tree (e.g. Hasegawa, 2017). The MgCO_3_ wt % is categorized into low magnesian calcite (LMC: 0–4 wt %), intermediate magnesian calcite (IMC: 4–8 wt %), and high magnesian calcite (HMC: >8 wt %). As highlighted in a blue zone, the MgCO_3_ data exhibit broad trends mostly consistent with the general phylogenetic evolution. Photosynthetic or photosymbiotic species are indicated as yellow data points, appearing in Rhodophyta, Foraminifera and Cnidaria; the two photosymbiotic species of Anthozoa are *Isis hippuris* and *Tubipora* sp. (Table 1). Red data points of Echinoidea correspond to specific skeletal parts of 3 regular species, indicating that their teeth and spines have significantly lower MgCO_3_ wt % (IMC) than the other skeletal parts (HMC) (Table 1).

The observed inter-taxonomic differences in (i) mineral phase and (ii) calcite-phase MgCO_3_ wt % are generally in agreement with previous studies (e.g. Chave, 1954; Thompson and Chow, 1955; Lowenstam, 1961; Kennedy et al., 1969; Rucker and Carver, 1969; Mackenzie et al., 1983; Lowenstam and Weiner, 1989) and must be as a result of the evolution/diversification of biomineralization processes according to the drastic changes in marine environments through the Phanerozoic eon (see the Introduction section). Our mineral-phase and MgCO_3_ data are likely to be more or less affected by environmental differences between sample sites. On the other hand, many species, belonging to different phyla/classes, were collected from one location (e.g. Wakayama) or some locations of similar environmental conditions (see Fig. 1 and annual-mean SST data of the sample locations listed in Table 2); therefore, the effect of environmental differences on our mineral-phase and MgCO_3_ data would be evidently small compared with the observed inter-taxonomic differences.

### Rhodophyta (Florideophyceae)

The Florideophyceae samples are calcareous red algae and consist of (i) calcite for 2 species, (ii) aragonite for 1 species and (iii) calcite-aragonite mixture for 3 species. The calcite-aragonite mixture species consist mainly of calcite with a trace or significant amount of aragonite. Of the 6 species analyzed, the aragonite species *Tricleocarpa cylindrica* belongs to the order Nemaliale, and the other 5 species to the order Corallinales; these results are in good agreement with a data compilation of Smith et al. (2012). From the viewpoint of their morphology, these calcifying algal species are categorized into two types as follows: (i) the geniculate (articulated) type and (ii) the non-geniculate (non-articulated) type. Applying the categorization to our results, the latter type corresponds to the calcite-aragonite mixture species, and the former type to the other species. Medaković et al. (1995) observed for some taxa of Corallinales as follows: (i) geniculate-species skeletons were composed of almost entirely calcite (e.g. *Amphiroa, Jania* and *Corallina*); (ii) non-geniculate-species skeletons were of predominantly calcite but frequently accompanied by a small amount of aragonite (e.g. *Lithophyllum* and *Phymatolithon*) and, in some cases, comprised of mainly aragonite with a significant amount of calcite (e.g. *Pseudolithophyllum*). These observations are mostly consistent with our results and Smith et al. (2012). Calcite phases of the 5 Corallinales species we analyzed are HMC (11.7–14.8 wt % MgCO_3_: Table 1 and Fig. 2), which is in good agreement with Smith et al. (2012) who reported that MgCO_3_ percentages of *Amphiroa, Corallina* and *Lithophyllum* species (collected around northern New Zealand: 29.2°S–38.3°S) were 13–16, 11–15 and 12–16 wt %, respectively; the latitude range is similar to that of the 5 species we collected in the northern hemisphere (i.e. ∼24°N–34°N).

### Foraminifera (Globothalamea and Tubothalamea)

The Globothalamea and Tubothalamea samples are HMC (11.0–11.7 wt % MgCO_3_: Table 1 and Fig. 2). Foraminifera are generally categorized into planktonic, benthic and encrusting types; some have photosynthetic symbionts inside of their cytoplasm and live in the surface ocean (i.e. the euphotic zone). The species analyzed in this study are benthic (*Baculogypsina sphaerulata, Calcarina* sp., *Amphisorus hemprichii* and *Marginopora* sp.) and encrusting (*Homotrema rubrum* and *Miniacina miniacea*); the 4 benthic species are well known to usually have photosynthetic symbionts. As for the 3 benthic species except *Marginopora* sp., MgCO_3_ data of our samples collected from Ishigaki and Okinawa Islands (annual-mean SSTs of ∼25–26°C: Table 2) are generally in good agreement with recently-reported data for samples grown at ∼24.5–27.5°C (Gacutan et al., 2017; Not et al., 2017; Singh, 2021), which seems to reinforce that our MgCO_3_ measurement is accurate. The recently-reported data are expressed as the Mg/Ca ratio, ranging from ∼140 to ∼170 mmol/mol, which is equivalent to the MgCO_3_ percentage from ∼10.5 to ∼12.5 wt %.

At least for benthic Foraminifera, there is a relatively clear tendency that photosymbiotic (non-photosymbiotic) species have HMC (LMC) shells (see a compilation table in Not et al., 2017), which is consistent with our data. Combining with the HMC data we obtained for (i) Rhodophyta species (see above) and (ii) two photosymbiotic Cnidaria species (see below) leads to a speculation that photosynthetic activities within calcite-secreting marine species tend to significantly enhance the Mg uptake into their calcitic shells/skeletons (see Fig. 2).

### Cnidaria (Anthozoa and Hydrozoa)

The Anthozoa samples are composed of (i) aragonite for 12 species, (ii) calcite for 8 species (9 individuals) and (iii) hard organic substance for 1 species, although two of the calcite species have, in part, hard organic skeletons (Table 1). All of the 8 calcite species belong to the order Alcyonacea of the subclass Octocorallia and are described as follows: (i) *Corallium japonicum* is known as a red precious coral, living in water depths of ∼100–350 m (Tsounis et al., 2010; Nonaka et al., 2012); (ii) *Dendronephthya aurea* is a soft coral, usually living in low-light conditions in a water-depth range from several meters to a few tens of meters, and has very minute calcitic spicules within its body; (iii) the genus *Isidella* is a kind of bamboo coral, living in water depths of several hundred meters, with its skeleton made up of calcitic internodes and hard-organic nodes (composed of gorgonin protein) (e.g. Ehrlich et al., 2006; Noé and Dullo, 2006; Ehrlich, 2019); (iv) *Isis hippuris* is also a kind of bamboo coral, usually living in tropical/subtropical shallow waters in the Indo-western Pacific region, with its skeleton constituted of calcitic internodes and hard-organic nodes (composed of gorgonin protein) (e.g. Rowley et al., 2015; Ehrlich, 2019; Saucier et al., 2021); (v) *Melithaea japonica* is a kind of sea fan, usually living in low-light conditions in a water-depth range from a few tens to several tens of meters (observed in Japan), with its skeleton made up of very minute calcitic spicules consolidated around thin organic axes (e.g. Matsumoto and van Ofwegen, 2015); (vi) *Pleurocorallium elatius* is known as a pink precious coral, living in water depths of ∼100–350 m; (vii) *P. konojoi* is known as a white precious coral, living in water depths of ∼100–350 m (Tsounis et al., 2010; Nonaka et al., 2012) and (viii) the genus *Tubipora* is commonly referred to as the red organ pipe coral (because of its skeletal color and shape), living in tropical/subtropical surface waters and growing into colonies usually a few tens to several tens of centimeters in size, rarely to be the dominant contributor to coral-reef formation (e.g. Richards et al., 2013). It is known that *I. hippuris* and *Tubipora* species, living in high-light conditions, have unicellular symbiotic algae (zooxanthellae) in their living tissues (thus termed ‘zooxanthellate species’), while that the other 6 calcite species, living in low- or no-light conditions, do not have the symbiotic algae (thus termed ‘azooxanthellate species’). The two zooxanthellate samples are HMC (9.5 and 12.5 wt % MgCO_3_: Table 1 and Fig. 2), supporting the above-mentioned speculation that photosynthetic activities within calcite-secreting marine species tend to significantly enhance the Mg uptake into their calcitic shells/skeletons. On the other hand, similar or higher MgCO_3_ values (11.5 and 13.3 wt %) are observed for the azooxanthellate samples *D. aurea* and *M. japonica*; the other 5 azooxanthellate samples have distinctly lower MgCO_3_ values (5.5–8.4 wt %). Both *D. aurea* and *M. japonica* secrete very minute spicules (see above and Table 1), with their bodies being very soft or flexible, whereas the other azooxanthellate calcite species secrete distinctly larger skeletons, with their bodies being very hard. The former species may significantly differ in biomineralization process from the latter species, which might cause the higher MgCO_3_ values of the former species. Our observations for *Isidella* sp. and *Isis hippuris* indicate that even the organic nodes contain a detectable amount of calcite (a few wt %). Of the 12 aragonite species, only *Heliopora coerulea* (known as the blue coral) belongs to the order Helioporacea of the subclass Octocorallia; the other 11 species belong to the order Scleractinia of the subclass Hexacorallia. The species *H. coerulea* is zooxanthellate and grows into a large colony (in some cases, more than several meters wide) in tropical surface oceans to be a significant contributor to coral-reef formation. From the viewpoint of their nature as significant coral-reef builders, the genera *Heliopora* and *Tubipora* are very unique within the subclass Octocorallia. Of the 11 aragonitic scleractinian hexacorals, *Cycloseris fragilis, C. tenuis, Dipsastraea speciosa*, two *Montipora* species, *Porites lutea*, and *P. australiensis* are zooxanthellate, whereas the other 4 species are azooxanthellate. The zooxanthellate genus *Cycloseris* is a solitary disc coral and does not significantly contribute to coral-reef formation, whereas the other 3 zooxanthellate genera are important coral-reef builders. Of the 4 azooxanthellate species, *Flabellum deludens* and *Stephanocyathus spiniger* are deep-sea solitary corals mostly found in water depths of ∼100 to ∼1000 m, while *Cyathelia axillaris* and *Dendrophyllia arbuscula* are colonial corals living in a water-depth range from a few meters to a few hundred meters and less contribute to coral-reef formation. The hard organic-skeleton species *Antipathes grandiflora* is a kind of black coral (or antipatharian coral) and belongs to the order Antipatharia of the subclass Hexacorallia, usually living in water depths of a few tens to a few hundreds of meters; the skeleton is composed primarily of chitin and antipathin protein (e.g. Ehrlich, 2019). Antipatharian corals had been considered to be azooxanthellate until Wagner et al. (2011) showed evidence, by using molecular typing and histology, that many of antipatharian species (including *Antipathes* species) in Hawaiian Islands and Johnston Atoll contain zooxanthellae as a symbiont.

The Hydrozoa samples are skeletons of aragonite, with a few to several wt % of calcite detected for one of the two *Millepora intricata* samples; the other *M. intricata, M. exaesa* and *M. tenella* samples consist of aragonite only (Table 1). The small amount of calcite detected may be due to some small calcite-secreting parasite in porous *Millepora* skeletons. It is known that some species (a few to several millimeters in size) of barnacles (e.g. *Pyrgoma* sp. of Cirripedia) and polychaete worms (e.g. *Phyllochaetopterus* sp. of Polychaeta) can be parasitic on *Millepora* skeletons (Lewis, 1989). As shown in Table 1, all the analyzed Cirripedia (barnacle) species have calcite skeletons, and all the analyzed Polychaeta species secrete tube shells of calcite-aragonite mixture. The polychaete-worm parasitism can be found also in the skeleton of scleractinian hexacorals of Anthozoa (most typically, *Spirobranchus giganteus* on *Porites* corals). The genus *Millepora* is zooxanthellate, living in tropical surface waters as reef builders, and is similar in appearance to scleractinian hexacorals; thus, although not a true coral, it is commonly referred to as the fire coral (because skin contact with it will result in immediate burning pain). *Stylaster profundiporus* is found in water depths of several meters to a few hundreds of meters; the genus *Stylaster* is azooxanthellate.

### Bryozoa (Gymnolaemata)

The Bryozoa samples belong to the class Gymnolaemata and consist of (i) aragonite for 1 genus (3 individuals of *Adeonellopsis* sp.), (ii) calcite for 19 species (23 individuals) and (iii) mixture of calcite with a trace amount (∼2–5 wt %) of aragonite for 2 species (*Alcyonidium foliaceum* and *Iodictyum phoeniceum*) (Table 1); *A. foliaceum* belongs to the order Ctenostomatida, and all of the other analyzed species belong to the order Cheilostomatida. The aragonite phase of *Adeonellopsis* is inconsistent with previous reports of this genus that indicate trace-to-small amounts (1–11 wt %) of calcite coexisting with aragonite (i.e. bimineralic phase) (e.g. Smith et al., 2006; Wejnert and Smith, 2008); the reason for this inconsistency is unclear. Taylor et al. (2006) reported that *Adeonella lichenoides* skeleton from Malaysia (5–6°N) was composed of aragonite only; *Adeonella* and *Adeonellopsis* belong to the same family Adeonidae. On the other hand, Smith and Clark (2010) found mineral phases of aragonite-calcite mixture or calcite for 24 specimens of *Adeonella* collected from southwestern Chile (42–52°S). The calcite single phases we observed for the genus *Biflustra, Bugula, Caberea, Celleporina, Iodictyum, Jellyella, Reteporella*, and *Thalamoporella* agree with previous studies (e.g. Smith et al., 1998, 2006; Hall et al., 2002; Kuklinski and Taylor, 2009; Krzeminska et al., 2016; Taylor et al., 2016; Loxton et al., 2018), with a possible exception of *I. phoeniceum* in which we detected a trace amount of aragonite. According to published data on the mineralogy of Cheilostomatida species, the calcite single phase is much more frequently observed than the calcite-aragonite mixed phase (bimineralic phase) or aragonite single phase in temperate to polar oceans, whereas the bimineralic phase is found for ∼10–35 % of specimens in tropical to temperate oceans (Poluzzi and Sartori, 1975; Borisenko and Gontar, 1991; Smith et al., 1998; Kuklinski and Taylor, 2009; Smith and Clark, 2010; Taylor et al., 2016; Loxton et al., 2018; Piwoni-Piórewicz et al., 2020). The former observation is consistent with our results (i.e. mostly the calcite single phase) obtained mostly from Wakayama located in a temperate region of Japan (∼33.5°N: see Fig. 1), although this site is evidently influenced by the warm Kuroshio Current originating from the northwestern tropical Pacific; the latter observation seems to be incompatible with our results (i.e. no distinct bimineralic phases), although the number of Cheilostomatida individuals (species) we analyzed is only 27 (20–22). The Ctenostomatida sample *A. foliaceum* has a MgCO_3_ percentage of 11.3 wt %, which is distinctly larger than the values of the other Cheilostomatida samples ranging from 1.9 to 9.3 wt % (Table 1). The wide MgCO_3_ range (from LMC to HMC) of Cheilostomatida is in good agreement with previous studies performed mostly in temperate to polar regions (e.g. Smith et al., 1998, 2006; Kuklinski and Taylor, 2009; Smith and Clark, 2010; Loxton et al., 2018; Piwoni-Piórewicz et al., 2020).

### Brachiopoda (Rhynchonellata and Lingulata)

Rhynchonellata species live mostly in very calm water conditions and are found most frequently in water depths from several tens to a few hundreds of meters (i.e. low- or no-light conditions), as our samples were collected from (see Table 1). All of the samples are LMC (1.0–2.1 wt % MgCO_3_: Table 1 and Fig. 2); these values are generally consistent with recent reports for Rhynchonellata shells, including *Laqueus rubellus, Terebratalia* spp. and *Terebratulina* spp. (Pérez-Huerta et al., 2008; Brand et al., 2013; Butler et al., 2015). These reports provide the following observations: (i) the shell Mg/Ca ratio varies from ∼2–7 mmol/mol at ∼6°C to ∼20–30 mmol/mol at ∼20°C, which is equivalent to a MgCO_3_ variation from ∼0.2–0.6 wt % at ∼6°C to ∼1.7–2.5 wt % at ∼20°C; (ii) the Mg/Ca ratio is significantly affected by some biological factors, including inter-genus differences; (iii) the outer layer and the posterior (juvenile) part of the shell have abnormally high Mg/Ca ratios.

Lingulata species inhabit vertical burrows in sandy/muddy sediments mostly in coastal shallow waters, with their habitat areas very restricted. For our sample *Lingula anatina*, we applied 3 different pretreatment methods to its shell (see the Materials and Methods section) and observed no significant differences in the XRD profile between the methods, all indicating the phase of fluorapatite[Ca_5_(PO_4_)_3_F]–hydroxyapatite[Ca_5_(PO_4_)_3_OH] solid solution (Table 1). This observation is consistent with previous reports (e.g. Kelly et al., 1965; Iijima and Moriwaki, 1990; Williams et al., 1994).

### Mollusca (Bivalvia, Cephalopoda, Gastropoda and Polyplacophora)

The Bivalvia samples are composed of (i) aragonite for 14 species and (ii) calcite-aragonite mixture for 12 species (14 individuals) (Table 1). Of the calcite-aragonite mixture species, *Crassostrea gigas, C. nippona* and *Mizuhopecten yessoensis* do not have any nacreous portions, whereas the other 9 species have nacreous portions as an inner layer of their shells. It is well known that any nacreous portions consist of aragonite. The genus *Crassostrea* is commonly referred to as true oysters and belongs to the order Ostreida; the genus *Mizuhopecten* is a kind of scallop and belongs to the order Pectinida. For the 3 non-nacreous species, we observed that their shells were predominantly composed of calcite, with a trace or significant amount of aragonite detected in two specific portions as follows: (i) the adductor muscle scar and (ii) the hinge part. As an exception, no aragonite was detected in the hinge part of one of the two *C. gigas* samples; the reason of this is unclear. Apart from this exception, our observations are almost consistent with previous studies for the *Crassostrea* and *Mizuhopecten* species (e.g. Stenzel, 1962, 1963; Carriker and Palmer, 1979; Chinzei, 1982; Lee et al., 2011; Zhao et al., 2015). The adductor muscle scar corresponds to an area where a thin combined layer of prismatic aragonite and organic matrix (named ‘myostracum layer’, roughly a few tens of μm in thickness for *Crassostrea*) is exposed directly to the inner surface of the shell; this muscle scar is surrounded by calcite materials (e.g. foliated and chalk layers in the case of *Crassostrea*) (Lee et al., 2011). All of the analyzed hinge parts of the 3 non-nacreous species include a resilifer, which is a pit or groove where the internal ligament (named ‘resilium’ composed of organic materials and aragonite) is located; however, we removed the resilium prior to XRD analysis, suggesting that another aragonitic part exists around the resilifer. Carriker and Palmer (1979) found a trace or significant amount of aragonite layered in the calcitic umbonal portion of 3 *Crassostrea* species and 3 *Ostrea* species (the resilium had been removed); this portion is probably equivalent to the hinge part we analyzed in the current study. For the mussel species *Mytilus galloprovincialis* belonging to the order Mytilida, we found significant variability in the calcite/aragonite percentage between the shell parts: (i) in the posterior part including adductor muscle scar, calcite (aragonite) is the major (minor) phase; in the central part, the opposite occurs; (iii) in the anterior part adjacent to umbo, aragonite overwhelms calcite. These observations are consistent with previous studies on calcite-aragonite distributions in *Mytilus* shells, including *M. galloprovincialis* and *M. edulis* (e.g. Taylor et al., 1969; Checa et al., 2007; Dalbeck, 2008; Hahn et al., 2012; Gao et al., 2015; Wan et al., 2019); they report that the shells are comprised of two main layers as follows: (i) an outer layer of prismatic calcite with the outermost thin organic layer termed ‘periostracum’ and (ii) an inner layer of nacreous aragonite with a thin layer of aragonitic myostracum. The former is secreted at the posterior edge for extending the shell length/height, while the latter is secreted on the inner surface of the former almost evenly and continuously, resulting in an increase, from the posterior via central to anterior part, of the thickness ratio of the aragonite to calcite layers (Milano et al., 2020); this increase of the aragonite/calcite ratio is probably what we observed in the *M. galloprovincialis* sample. All of the other 8 calcite-aragonite mixture species belong to the order Ostreida: *Atrina pectinata japonica, A. vexillum, Electroma alacorvi, Isognomon isognomum, Malleus albus* (hammer oyster), *Pinctada fucata martensii* (Akoya pearl oyster), *P. margaritifera* (black-lip pearl oyster), and *P. maxima* (South Sea pearl oyster); we found that these species have an outer prismatic layer and an inner nacreous layer composed of calcite and aragonite, respectively; these observations are consistent with general knowledge of biomineralogy of these species (e.g. Kennedy et al., 1969; Taylor et al., 1969). Only LMC phases were observed for the 12 species of calcite-aragonite mixture (∼0.0–3.3 wt % MgCO_3_: Table 1 and Fig. 2). As for the 14 species of aragonite single phase, we observe that only *Septifer bilocularis* has a nacreous portion as an inner layer, with a non-nacreous outer layer, and that the other 13 species have no nacreous portions but are entirely porcellaneous. The observation for *S. bilocularis* is consistent with Owada and Hoeksema (2011) but not with Taylor et al. (1969) who reported an outer nacreous layer and an inner prismatic layer. Although the two genera *Septifer* and *Mytilus* belong to the same family Mytilidae, the shell mineralogy of *S. bilocularis* (entirely aragonite) is in contrast to that of *Mytilus* species (calcite-aragonite mixture). Of all the Bivalvia species analyzed in this study, only *Tridacna maxima* (small giant clam) contains algal symbionts (zooxanthellae), with its shell composed of aragonite; these features are the same as those of scleractinian reef-building hexacorals. The genus *Tridacna* inhabits tropical/subtropical coral reefs in the Indo-Pacific region.

The Cephalopoda samples consist of (i) calcite for 2 *Argonauta* species and (ii) aragonite for 1 *Sepia* species, which is consistent with previous studies (e.g. Kelly, 1901; Bøggild, 1930; Lowenstam and Weiner, 1989); MgCO_3_ percentages of the *Argonauta* samples are 5.4 and 6.7 wt % (i.e. IMC) (Table 1 and Fig. 2). The genera *Argonauta* and *Sepia* belong to the same subclass Coleoidea but to different orders Octopoda and Sepiida, respectively. The *Argonauta* shell is secreted only by the female and used for an eggcase, whereas the *Sepia* cuttlebone is formed in both the male and female and used for buoyancy control. A well-known shell-bearing family of Cephalopoda is Nautilidae, which belongs to the subclass Nautiloidea; shells of Nautilidae species consist of prismatic and nacreous aragonite layers (e.g. Bøggild, 1930; Lowenstam and Weiner, 1989). It is well known that the Nautilidae shell is divided into chambers with the animal only occupying the largest, outermost chamber; the other chambers are used for buoyancy control and are developmentally and functionally homologous to the aragonitic cuttlebone of Sepiida.

The Gastropoda samples are composed of (i) aragonite for 9 species and (ii) aragonite-calcite mixture for 8 species, with the latter species having LMC phases (0.6–3.8 wt % MgCO_3_); similar features can be seen in the Bivalvia samples (Table 1 and Fig. 2). The *Janthina globosa* and *J. janthina* samples have distinctly higher MgCO_3_ values (3.5–3.8 wt %) than the other species of aragonite-calcite mixture (0.6–2.6 wt %). The genus *Janthina* are pelagic and planktonic sea snails living in tropical/subtropical oceans, with their shells usually growing up to ∼3–4 cm in size; they secrete transparent mucous that hardens into a bubble raft which keeps them floating at the oceans surface, and feed on some kinds of small jellyfishes and crustaceans. This living mode is unique among Gastropoda species and might contribute to the higher MgCO_3_ values. The *Janthina* samples were collected at Okinawa and Ishigaki Islands, where annual-mean SSTs are ∼25–26°C (Table 2); however, they are likely to have been conveyed via the warm Kuroshio Current from the northwestern tropical Pacific where SST is presumably ∼2–3°C higher than that around Okinawa/Ishigaki Islands. This is probably the most likely explanation for the higher MgCO_3_ values of the *Janthina* samples.

For the *Cellana testudinaria* shell with no nacreous portions, a significant amount of calcite is detected in both the inner layer and outer thin layer, where the calcite-to-aragonite ratio is larger in the outer thin layer. For the *Haliotis madaka* and *Turbo sazae* shells with an inner nacreous layer, a significant amount of calcite is detected only in the outer thin layer, with the other shell parts composed of aragonite only. The calcite in the outer thin layer is probably secreted immediately under the periostracum (the outermost very thin organic layer), as observed in some bivalves (e.g. the genera *Mytilus* and *Pinctada*: see above). It has been observed that *Haliotis* shells have, immediately under the periostracum, an outer thin prismatic layer composed of calcite, aragonite or both, depending on species (e.g. Nakahara et al., 1982; Mutvei et al., 1985; Dauphin et al., 1989, 2014; Cusack et al., 2013); for example, the outer prismatic layer of *H. rufescence* is composed of calcite, whereas those of *H. asinina* and *H. gigantea* are of aragonite. Similarly to the *H. madaka* and *T. sazae* shells, the *Guildfordia triumphans* and *Omphalius pfeifferi carpenteri* shells have an inner nacreous and outer non-nacreous layer; the whole-shell XRD analyses indicate aragonite single phase for *G. triumphans* and aragonite-calcite mixture for *O. pfeifferi carpenteri*, suggesting that the outer non-nacreous layers of the former and latter species consist of aragonite and calcite (or calcite-aragonite mixture), respectively. Mannino (2000) and García-Escárzaga et al. (2015) observed an inner nacreous layer and outer calcite layers for shells of *Phorcus lineatus*, which, together with *T. sazae* and *O. pfeifferi carpenteri*, belongs to the superfamily Trochoidea. The whole-shell XRD analysis of *Nerita incerta* suggests non-nacreous aragonite and calcite layers, which is compatible with Nehrke and Nouet (2011) who found that a *N. undata* shell was composed of an inner non-nacreous aragonite layer and an outer calcite layer.

All the Polyplacophora samples are composed of aragonite with respect to shell plates; in addition, girdle spines of *Acanthopleura gemmata* and *Liolophura japonica* are also composed of aragonite (Table 1). All the 7 species analyzed belong to the order Chitonida; *Acanthochitona achates* and *Stenoplax alata* belong to the families Acanthochitonidae and Ischnochitonidae, respectively, while the other 5 species to the family Chitonidae. Our results are compatible with previous observations for Chitonida species as follows: (i) multi-layered structures with varying types of aragonite (e.g. granular, crossed-lamellar, prismatic) in their shell plates (e.g. Haas, 1976; Laghi and Russo, 1978; Carter and Hall, 1990; Connors et al., 2012) and (ii) prismatic aragonite structures in their girdle spines (Treves et al., 2003).

### Annelida (Polychaeta)

The Polychaeta samples (*Hydroides* and *Spirobranchus*) belong to the family Serpulidae and exhibit similar mineral compositions of ∼85–65 wt % calcite and ∼15–35 wt % aragonite (Table 1). Although we have analyzed only 4 species (7 individuals) of the 2 genera, our results do not seem to be consistent with previous reports on the mineralogy of Serpulidae shells that indicate extremely large variability of the calcite (aragonite) percentage from ∼2 to ∼100 wt % (∼98 to ∼0 wt %) (Vinn et al., 2008; Smith et al., 2013; Ippolitov and Rzhavsky, 2015). For example, Vinn et al. (2008) observed mineral compositions of 98 wt % calcite and 2 wt % aragonite for *Hydroides norvegicus* and 5 wt % calcite and 95 wt % aragonite for *H. spongicola*, indicating an extreme difference within the same genus. Reasons for the difference between our results and previous reports and for the extreme inter-species difference of *Hydroides* are unclear but might be due to some biological and/or environmental difference. The calcite phases of our 7 samples are HMC (9.7–12.8 wt % MgCO_3_) (Table 1 and Fig. 2), suggesting that *Hydroides* may have slightly higher MgCO_3_ contents than *Spirobranchus* in the same environmental condition (see the data from Wakayama and Yamaguchi).

### Arthropoda (Cirripedia)

The Cirripedia samples are LMC (0.1–3.5 wt % MgCO_3_) except for one sample of *Capitulum mitella* (4.3 wt % MgCO_3_) (Table 1 and Fig. 2); the other samples of *C. mitella* have MgCO_3_ values of 2.3–3.5 wt %, which are significantly higher than those of all the other 6 species (0.1–1.2 wt %). The *C. mitella* samples were collected in intertidal zones in Wakayama and Ehime, where the samples *Megabalanus rosa* (0.2 wt % MgCO_3_), *M. volcano* (1.0 wt % MgCO_3_) and *Balanus trigonus* (0.9 wt % MgCO_3_) were also collected at water depths of 0–5 m or 5–10 m. The higher MgCO_3_ values of *C. mitella* may be related to the facts that *C. mitella* belongs to the order Pollicipedomorpha and that the other 6 species belong to the order Balanomorpha, and might reflect significantly different biomineralization processes between the two orders. The MgCO_3_ data for each specific skeletal part of *Balanus rostratus, C. mitella*, and *Striatobalanus tenuis* show within-individual variations of 0.1–1.0, 2.3–3.5, and 0.4–1.0 wt %, respectively. Except for the tergum of *C. mitella* (3.5 wt %), each of the within-individual variations is approximately comparable to the MgCO_3_ measurement reproducibility (≤ 0.5 wt %: see the Materials and Methods section); these results might suggest that, for Cirripedia species, within-individual variability of biomineralization processes is generally not significant.

### Echinodermata (Asteroidea, Crinoidea and Echinoidea)

The Asteroidea and Crinoidea samples are HMC (12.3–14.3 and 11.1–12.2 wt % MgCO_3_, respectively), while the Echinoidea samples are IMC or HMC (5.0–12.3 wt % MgCO_3_) (Table 1 and Fig. 2); the higher values of the Asteroidea species may be related to some inter-taxonomic difference of biomineralization processes within the phylum Echinodermata. The 5 Asteroidea species belong to the 3 orders Valvatida (*Certonardoa semiregularis, Fromia monilis* and *Patiria pectinifera*), Paxillosida (*Astropecten polyacanthus*) and Forcipulatida (*Coscinasterias acutispina*). The 3 Crinoidea species belong to the orders Comatulida (*Anneissia japonica* and *Capillaster* sp.) and Isocrinida (*Metacrinus rotundus*); the latter taxon develops a long stalk, but the former taxon does not (Table 1). *M. rotundus* is found most frequently near the edge of the continental shelf off the western coast of Japan (in 100–200 m water depth) and is commonly referred to as the Japanese sea lily; this is the shallowest-living species among the extant stalked crinoids. As for the *M. rotundus* and *Capillaster* sp. samples, no significant differences of the MgCO_3_ content are found between specific skeletal parts (11.5–12.2 and 11.1–11.3 wt %, respectively: Table 1), which may suggest that within-individual variability of biomineralization processes is not significant for Crinoidea species.

In contrast, the 3 Echinoidea samples *Diadema setosum, Echinometra mathaei* and *Heliocidaris crassispina* indicate distinct MgCO_3_ differences between specific skeletal parts of each; their teeth and spines are IMC (5.0–7.3 wt %), whereas the other skeletal parts (test plates, hemipyramids, epiphyses and rotulae) are HMC (9.9–12.3 wt %) (Table 1 and Fig. 2). This finding may be related to the fact that the teeth and spines, in contrast to the other skeletal parts, are abraded or chipped off when used in predation and protection, suggesting that the lower MgCO_3_ content of the teeth and spines may be due to functional adaptations through the evolutionary selection; it has been generally observed for biogenic calcites that the lower the MgCO_3_ content, the lower the hardness and solubility (e.g. Morse and Mackenzie, 1990; Railsback, 2006; Kunitake et al., 2012; Long et al., 2014; see the Introduction section). These 3 species and the 2 species *Mesocentrotus nudus* and *Strongylocentrotus intermedius* analyzed for their test plates only are categorized as ‘regular echinoids’, while the other 5 species analyzed are as ‘irregular echinoids’ belonging to the infraclass Irregularia. The regular and irregular echinoids differ distinctly in morphology and living mode; the former, living usually on rocky/sandy sea beds or in rock crevices and feeding mainly on algae and kelps, has a round-formed body with well-developed spines and complex mouth-apparatus known as Aristotle’s lantern (composed of teeth, hemipyramids, epiphyses and rotulae: see Table 1), whereas the latter, usually burrowing under seafloor sediments and feeding on fine organic detritus, has a flattened or heart-shaped body with thin/short spines and a simply structured mouthpart. Our observation for the 3 regular echinoids that their teeth and spines have distinctly lower MgCO_3_ wt % than the other skeletal parts is in agreement with previous reports for some regular echinoids that spines have significantly lower MgCO_3_ wt % or Mg/Ca ratios than test plates (e.g. Chave, 1954; Ries, 2004). Thus, it may be suggested for regular echinoids that biomineralization processes of their teeth and spines are significantly different from those of the other skeletal parts; in brief, within-individual variability of biomineralization processes may be significant for regular echinoids. Test plates of the *M. nudus* and *S. intermedius* samples have remarkably lower MgCO_3_ values (5.0–6.0 wt %, IMC) than those of the other regular echinoid samples (11.1–12.3 wt %, HMC) (Table 1 and Fig. 2); this is because the former samples grew under significantly lower SSTs in Hokkaido (Table 2: see discussion in the following subsection). Tests of the irregular echinoid *Scaphechinus mirabilis* (4 individuals from different locations) have distinctly lower MgCO_3_ values (6.9–7.5 wt %, IMC) than those of the other irregular species (9.5–12.0 wt %, HMC) (Table 1 and Fig. 2); this is not due to seawater-temperature differences but presumably to inter-taxonomic differences within Echinoidea.

### Assessment of the correlation of the calcite-phase MgCO_3_ data with seawater temperature

We assess the calcite-phase MgCO_3_ data obtained in this study, with respect to the well-known positive correlation between the Mg (or MgCO_3_) content of calcite and water temperature under which the calcite is formed inorganically or biogenically (see Introduction section). Fig. 3 shows scatter plots between our MgCO_3_ data for near-sea-surface species (≤ 30 m in water depth: see Table 1) and annual-mean SST data of their habitat regions (Table 2); these plots are given for all the biological classes analyzed for calcite-phase MgCO_3_ wt % except for the Rhynchonellata and Cephalopoda species that inhabited in deeper, wide-ranging, or unknown water depths (see Table 1). Pearson’s correlation analysis (the least-squares linear regression) is performed in each plot with the number of data points ≥ 5. High or moderate positive correlations between the MgCO_3_ and SST data are found for Bivalvia (*n* = 26, *r*^*2*^ = 0.63, *p* < 0.0001, SST range = 19.1°C), Gastropoda (*n* = 9, *r*^*2*^ = 0.91, *p* < 0.0001, SST range = 9.4°C) and Echinoidea (*n* = 19, *r*^*2*^ = 0.88, *p* < 0.0001, SST range = 18.6°C) (Figs. 3e–f and 3k), where (i) data points of *C. testudinaria, S. mirabilis* and regular echinoids’ teeth/spines are regarded as outliers and excluded, and (ii) SST data for the *J. globosa* and *J. janthina* samples are assumed to be 28°C (see the above discussion on the Gastropoda samples); selecting the data points of test plates of 5 regular-echinoid species gives a very high correlation (*n* = 6, *r*^*2*^ = 0.99, *p* < 0.0001: Fig. 3k), suggesting little effect of inter-taxonomic differences on the Mg–temperature relationship of regular-echinoid test plates. For the other 8 biological classes, the MgCO_3_–SST correlations are not significant (*p* > 0.05). In the plot of Cirripedia (Fig. 3h), data points of *C. mitella* appear to be outliers, where the MgCO_3_–SST correlation is not significant whether or not the *C. mitella* data points are excluded. Our inference for *C. mitella* and *S. mirabilis* that their anomalous MgCO_3_ values are presumably or possibly due to inter-taxonomic differences (see the above discussions on the Cirripedia and Echinoidea samples) seems to be reinforced by the MgCO_3_ vs. SST plots.

**Fig. 3.**
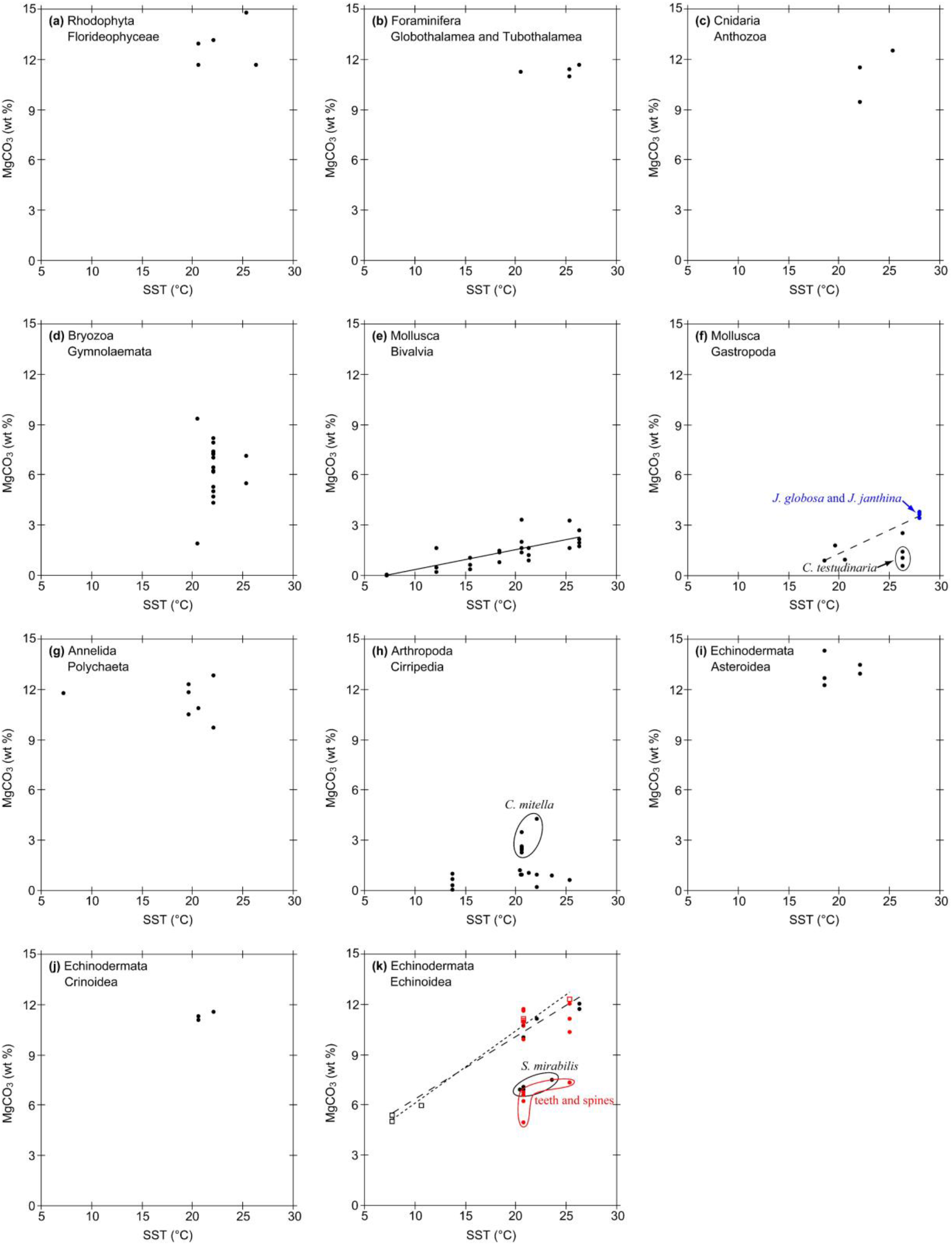
Scatter plots of the calcite-phase MgCO_3_ wt % of shells/skeletons of extant near-sea-surface species (≤ 30 m in water depth) collected in Japan versus annual-mean SSTs (°C) of their habitat regions: **(a)** Florideophyceae, Rhodophyta; **(b)** Globothalamea and Tubothalamea, Foraminifera; **(c)** Anthozoa, Cnidaria; **(d)** Gymnolaemata, Bryozoa; **(e)** Bivalvia and **(f)** Gastropoda, Mollusca; **(g)** Polychaeta, Annelida; **(h)** Cirripedia, Arthropoda; **(i)** Asteroidea, **(j)** Crinoidea and **(k)** Echinoidea, Echinodermata. These plots are drawn using the data shown in Tables 1 and 2. As indicated by ellipses, data points of *C. testudinaria, C. mitella* and *S. mirabilis* may be regarded as outliers. In the plot of Echinoidea, red data points represent specific skeletal parts of 3 regular species, indicating that their teeth and spines have distinctly lower MgCO_3_ values than the other skeletal parts (see Table 1 and Fig. 2). Pearson’s correlation analysis has been applied to each plot with the number of data points ≥ 5. The correlation analysis gives high or moderate MgCO_3_–SST correlations for the Bivalvia, Gastropoda and Echinoidea data, where (i) data points of *C. testudinaria, S. mirabilis*, and regular echinoids’ teeth/spines are excluded and (ii) SST data for the *J. globosa* and *J. janthina* samples are assumed to be 28°C (see discussion on the Gastropoda samples); for all the other biological classes, the MgCO_3_–SST correlations are not significant (*p* > 0.05). The MgCO_3_–SST correlations for Bivalvia, Gastropoda and Echinoidea are respectively as follows: *y* = –0.81 + 0.117*x* (*n* = 26, *r*^*2*^ = 0.63, *p* < 0.0001 ; straight line), *y* = –4.28 + 0.280*x* (*n* = 9, *r*^*2*^ = 0.91, *p* < 0.0001; dashed line) and *y* = 2.73 + 0.368*x* (*n* = 19, *r*^*2*^ = 0.88, *p* < 0.0001; dashed line). Selecting the data points of test plates of 5 regular Echinoidea species (open squares) gives a very high correlation as follows: *y* = 1.80 + 0.432*x* (*n* = 6, *r*^*2*^ = 0.99, *p* < 0.0001 ; dotted line).

The scatter plots of the 8 biological classes with non-significant MgCO_3_–SST correlations are mostly characterized by smaller numbers of data points (*n* = 3, 5 or 7; except for the Gymnolaemata [*n* = 20] and Cirripedia [*n* = 18] classes: Figs. 3d and 3h) and narrower SST ranges (1.5–5.7°C; except for the Polychaeta [14.9°C] and Cirripedia [11.6°C] classes: Figs. 3g and 3h), compared with those of the 3 classes with significant positive MgCO_3_–SST correlations (see above for *n* values and SST ranges). Accordingly, for each of the 8 biological classes, increasing the number of data points and/or widening the SST ranges might allow us to find significant positive MgCO_3_–SST correlations and distinct outliers (due to inter-taxonomic differences) that should be excluded from the correlations, as found in the Echinoidea data (Fig. 3k); this strategy, however, may not be effective for the Cirripedia and Gymnolaemata classes that already have larger numbers of data points with SST ranges of 11.6°C and 4.8°C, respectively (Figs. 3d and 3h). Very few previous studies have reported positive Mg–temperature correlations for shells/skeletons of Cirripedia and Gymnolaemata (Bryozoa) (Chave, 1954).

Our study method based on the whole-shell/skeleton XRD analysis is not the best one for assessing the MgCO_3_–temperature correlation (i.e. the Mg content vs. temperature correlation). The best method would be as follows: (i) measuring the Mg concentration along the shell’s or skeleton’s growth line of a specific species (by means of micro sampling or micro analysis such as the laser-ablation inductively-coupled plasma mass spectrometry, i.e. LA-ICP-MS), which reveals temperature-induced seasonal variation of the Mg concentration, then (ii) comparing the seasonal Mg variation with seasonal seawater-temperature variation under which the shell/skeleton grew; thereby, the Mg–temperature correlation of the specific species are accurately assessed. This kind of method, in contrast to our method, can find or detect (i) specific portions with abnormally-high Mg concentrations that cannot be ascribed to temperature effect but presumably to some biological effect (such portions cannot be used for assessing the Mg–temperature correlation) as well as (ii) even smaller inter-taxonomic differences of the Mg–temperature relationship (e.g. Freitas et al., 2005; Pérez-Huerta et al., 2008; Butler et al., 2015). It is very likely for near-sea-surface species that the secretion rate of their shells/skeletons significantly varies seasonally; for example, some species may cease the secretion in winter. In this case, the MgCO_3_ value obtained by our method (i.e. the whole-shell/skeleton analysis) does not correspond to the annual-mean SST, resulting in a method-induced error in the MgCO_3_–SST correlation; in contrast, the micro-sampling/analysis method can reveal the seasonal variation of the shell/skeleton-secretion rate and give an accurate Mg–temperature correlation. The uncertainty of our XRD-based MgCO_3_ measurements (≤ 0.5 wt %: see the Materials and Methods section) is close to or equivalent to lower MgCO_3_ values (0–1.0 wt %) of the LMC species (e.g. Bivalvia, Gastropoda and Cirripedia: Figs. 3e–f and 3h), which presumably more or less obscures their MgCO_3_–SST correlations. Hence, these problems of our method disturb and lower the MgCO_3_–SST correlation; this effect would vary depending on taxa and their shells’/skeletons’ MgCO_3_ contents. The very high MgCO_3_–SST correlation found for the HMC test plates of 5 regular-echinoid species (Fig. 3k) suggests that effects of both (i) the methodological problems and (ii) inter-taxonomic differences of regular echinoids are negligible.

In summary, the XRD-based methodology employed in this study is effective to investigate inter-taxonomic differences in the mineral phase and MgCO_3_ content of calcareous marine-species shells/skeletons but is less useful for assessing the correlation between the MgCO_3_ content and temperature than the micro-sampling/analysis method. Nevertheless, further accumulation of the XRD-based MgCO_3_ data of calcite-containing shells/skeletons secreted under wide-ranging seawater temperatures would probably provide more accurate assessment of the MgCO_3_–temperature correlation, especially for the investigated biological classes with smaller numbers of the MgCO_3_ data.

## ACKNOWLEDGMENTS

This study was funded by the Ministry of Economy, Trade and Industry (METI), Japan as part of its R&D supporting program titled ‘Development of Evaluation Technology for Long-term Geosphere Stability on Geological Disposal Project of Radioactive Waste (Fiscal Years 2013–2017)’. We thank Drs. Hiroya Yamano and Masato Kiyomoto for their support in sample collection; this study is a contribution to a monitoring project for global warming effects on marine environment funded by the Center for Global Environmental Research, National Institute for Environmental Studies, Japan.

Lastly, the first author (T.M.) dedicates this article to his mother Kiyo Mitsuguchi, who gave him warm encouragement for this research and passed away on 4 April 2022 at the age of 89.

## COMPETING INTERESTS

All the authors declare that no competing interests exist.

## AUTHOR CONTRIBUTIONS

Sample collection and species identification by TM, KM, KS, MH and MAY; experimental design by all authors; conduct of experiments by TM and YSK; data discussion and interpretation by all authors; writing the manuscript by TM and YSK.

## SUPPLEMENTARY MATERIALS

There are no supplementary materials on this article.

## Notes

### Competing Interest Statement

The authors have declared no competing interest.

